# Molecular signature of human endometrial stem/progenitor cells at the single cell level

**DOI:** 10.1101/2025.06.23.660982

**Authors:** Harriet C. Fitzgerald, Sally Mortlock, Caitlin E. Filby, Fiona L. Cousins, Elizabeth Marquez-Garcia, Katherine A. Wyatt, Brett McKinnon, Luk Rombauts, Jim Tsaltas, Grant W. Montgomery, Caroline E Gargett

**Author notes:** Authors contributed equally (Harriet C. Fitzgerald, Sally Mortlock). The authors declare no competing interest.

## Abstract

Human endometrium sheds and regenerates each month during the menstrual cycle. N-cadherin^+^ (*CDH2*) glandular epithelial progenitors and SUSD2^+^ MSC and their niches have been identified but their signaling interactions remain unknown. SSEA1^+^ epithelial cells resurface the endometrium generating a new luminal epithelium each cycle. Based on these markers, we characterised the gene expression of human endometrial stem/progenitor cell-enriched populations derived from FACSorted hysterectomy endometrium by scRNAseq. Two of 10 epithelial clusters contained *CDH2^+^SOX9^+^* cells with high *TRH* and *IHH* expression. N-cadherin and IHH, and SSEA1 and Hedgehog coreceptor BOC immunocolocalised in the basal layer of endometrial glands from which the new functional layer glands regenerate each menstrual cycle. Two of six mesenchymal clusters contained *SUSD2^+^* MSC, one with high *MUSTN1* expression. Epithelial progenitors and endometrial MSC transitioned to their respective progeny. We provide new insights into human endometrial stem/progenitor cell signaling pathways and niche interactions regulating their function.

**Highlights:**

- Enriching cell suspensions of human endometrial cells from hysterectomy tissues with CDH2^+^ epithelial progenitors, SSEA-1^+^ basalis epithelial cells and SUSD2^+^ perivascular MSC enabled their gene profiling at the single cell level.
- Multiple cell fate trajectory analyses showed that *CDH2^+^SOX9^+^* epithelial cells and *SUSD2^+^MYH11^+^MCAM^+^MUSTN1^+^* MSC are the progenitors for human endometrial glandular and luminal epithelial cells, and stromal vascular cells, respectively.
- Enriched *TRH* expression in human endometrial epithelial progenitors suggests a role in their progenitor function.
- Interaction between IHH of SOX9^+^ human endometrial epithelial progenitors with basalis fibroblasts and decidualized stroma and co-receptor BOC suggests IHH signalling may have a role in basalis epithelial migration to repair the luminal epithelium during menses.

**eTOC blurb:** Gargett and colleagues provide a molecular signature of human endometrial stem/progenitor cells by enriching 4 stem/progenitor populations from hysterectomies using known markers for scRNAseq analysis. The two epithelial and two MSC progenitor states identified transitioned to their differentiated progeny. IHH was validated as a key signalling molecule of epithelial progenitors in basal glands that interacted with BOC co-receptors.

## Introduction

The endometrium (uterine lining) is the site where an embryo implants and placentation occur to support pregnancy. Human endometrium is one of the body’s most highly regenerative tissues. Each menstrual cycle, the upper two thirds (functionalis) is shed during menstruation, leaving a raw surface on the remaining basalis, which is rapidly re-epithelialized in the absence of the hormone estrogen (Gargett et al., 2012; Kaitu’u-Lino et al., 2010). As circulating estrogen levels rise during the proliferative stage of the cycle a new functionalis regenerates, with glands sprouting vertically from the horizontal interconnected glands of the basalis (Yamaguchi et al., 2021). Following ovulation, during the secretory phase of the cycle, ovarian progesterone induces gland differentiation that produce secretions (histotroph) which nourishes the embryo during the early weeks of placentation (Burton et al., 2002). In the absence of an implanting embryo, circulating progesterone falls and the functionalis is shed piecemeal into menstrual fluid. Up to one cm of endometrial tissue regenerates each month, comprising glands lined by a columnar epithelium and a vascularized stroma. More than 400 menstrual cycles occur in modern reproductive age women (Salamonsen et al., 2021).

The first evidence for human endometrial stem/progenitor cells was the demonstration of clonogenic epithelial and stromal cells derived from full-thickness (hysterectomy) endometrium (Chan et al., 2004). Single cells initiating large epithelial or stromal clones underwent self-renewal during serial cloning, generated ∼6×10^11^ cells. Epithelial clones differentiated into 3D gland-like structures in vitro (Gargett et al., 2009). Large stromal clones differentiated into multiple mesodermal lineages in vitro. This indicates that human endometrium contains epithelial progenitors and MSC-like cells.

Markers enriching for clonogenic endometrial MSC-like cells, co-expression of CD140b and CD146 (Schwab and Gargett, 2007), identified their perivascular niche in both the functionalis and basalis. A single perivascular marker, SUSD2, also enriched for clonogenic MSC and differentiate into stromal tissue and migrating endothelial cells in vivo (Masuda et al., 2012).

Endometrial epithelial progenitor markers were first identified by comparing purified epithelial cells from basalis-like postmenopausal to premenopausal endometrium by gene microarray (Nguyen et al., 2017). Atrophic menopausal endometrium responds like the basalis and regenerates a functionalis when exposed to menopausal estrogen therapy. *CDH2*, encoding N-cadherin was among 11 differentially expressed surface markers genes upregulated in post-menopausal endometrium (Nguyen et al., 2017). Greater epithelial progenitor activity was demonstrated in N-cadherin^+^ than N-cadherin^−^ epithelial cells. Their basalis location was shown in the horizontal branching glands adjacent to the myometrium. AXIN-2 and SSEA1 were discovered as basalis epithelial markers (Nguyen et al., 2012; Valentijn et al., 2013). SSEA1^+^ identified basalis epithelial cells with intermediate telomerase activity and longer telomeres, suggesting progenitor activity. SSEA1^+^ epithelial cells are nuclear SOX9^+^ and located at an ill-defined basalis-functionalis junction, where they may migrate from basalis gland stumps during menses to resurface the raw endometrium, explaining their extra location in the luminal epithelium (Cousins et al., 2021).

Recent single cell RNA sequencing (scRNAseq) studies identified multiple endometrial epithelial, stromal, and immune cell subpopulations in human endometrium (Fonseca et al., 2023; Garcia-Alonso et al., 2021; Lv et al., 2022; Marečková et al., 2024; Queckbörner et al., 2021; Tan et al., 2022; Wang et al., 2020). Most identified endometrial MSC (eMSC) as studies have mainly used endometrial biopsies, which sample the functionalis(Queckbörner *et al*., 2021; Tan *et al*., 2022; Wang *et al*., 2020)(Queckbörner *et al*., 2021; Tan *et al*., 2022; Wang *et al*., 2020). These found our three published eMSC markers, but N-cadherin^+^ epithelial progenitors were mostly not identified, likely due to their deep basalis location not sampled in a biopsy, although *AXIN2*^+^ cells were observed. Two clusters comprising many *SOX9*^+^ epithelial cells were found in functionalis glandular epithelial cells during the proliferative phase, 40-fold more than *SOX9^+^* basalis and luminal epithelial cells (Marečková *et al*., 2024). *SOX9*^+^ cells appear more numerous than expected for solely a basalis and luminal epithelial marker (Garcia-Alonso *et al*., 2021).

To gain insight into the stem/progenitor cell state of epithelial progenitors and eMSC and their respective niche cells that regulate their cell fate decisions, we enriched human endometrial stem/progenitor cells by FACSorting hysterectomy-derived endometrial single cell suspensions using our surface markers, N-cadherin, SSEA1 and SUSD2 and recombined them with their mature progeny for subsequent scRNAseq.

## Results

### Cell identities defined by scRNA seq in enriched human endometrial stem/progenitor cell populations

To generate a molecular signature of human endometrial stem/progenitor cells, cells were first isolated from hysterectomy endometrium, containing both basalis and functionalis from 5 women not taking hormones. Single cell suspensions were enriched for epithelial progenitor cells and eMSCs by FACS using known markers (Figure S1), generating 6 subpopulations of endometrial cells: 3 basalis epithelial progenitors; EpCAM^+^N-cadherin^+^SSEA1^−^, EpCAM^+^N-cadherin^+^SSEA1^+^ and EpCAM^+^N-cadherin^−^SSEA-1^+^; mature epithelial cells, EpCAM^+^N-cadherin^−^SSEA1^−^; eMSCs, SUSD2^+^EpCAM^−^ and stromal fibroblasts, SUSD2^−^EpCAM^−^. The 6 subpopulations were pooled to enrich for the stem/progenitor cells and underwent scRNAseq (Figure 1A). An average of 2,346 cells were sequenced from each sample with a median of 3,105 genes detected per cell (Table 1). Following sample integration, cell clustering was performed at 0.4 resolution based on cluster stability and identification of known biological groups and cell types (Figure 1B). Sixteen cell clusters were identified (Figure 1C). The number of cells within a cluster contributed by each subject varied (Figure S2A), likely due to the cycle stage when tissue was harvested and individual pathologies (Table S1). However, all samples contributed cells to each cluster (C).

**Figure 1.**
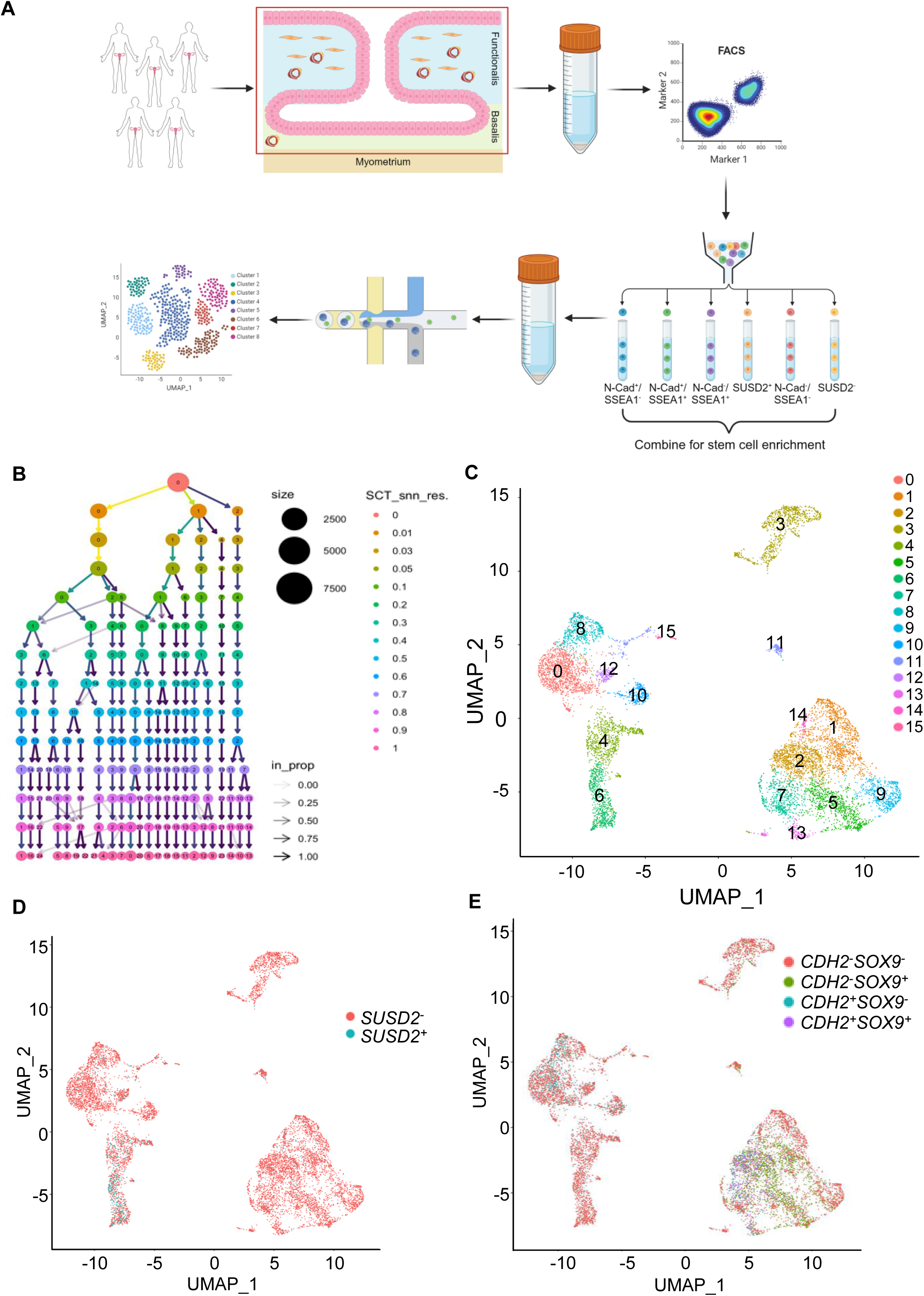
ScRNAseq analysis of human endometrial stem/progenitor cell-enriched endometrial cells. (**A**) Workflow from collection of (n=5) endometrial tissues from subjects undergoing hysterectomy, isolation of cells from the basalis and functionalis glands and stroma (red box, excluding myometrium), FACS sorted on published stem/progenitor cell markers, recombined in equal proportions where possible and analysed by scRNAseq. (**B**) Clustree visualisation of clusters generated by Seurat’s Louvain algorithm for varying resolution parameters (rows). Each circle represents a cluster identified at a given resolution, with size corresponding to the number of cells contained. Arrows between clusters illustrate cells transition across resolutions, with arrow shading showing the proportion of cells carried over from clusters at the preceding resolution. (**C**) UMAP showing 16 defined cell clusters at 0.4 resolution. (**D**) UMAP of *SUSD2^+^* eMSCs and *SUSD2^−^* cell populations. (**E**) UMAP of *CDH2* and *SOX9* expression in the cell populations. Clusters 2 and 7 contain higher numbers of *CDH2^+^* and *SOX9^+^* cells, likely endometrial epithelial progenitor populations.

**Table 1.** Single-cell RNA-seq metrics for each sample.

| Donor ID | Estimated Number of Cells | Mean Reads per Cell | Median Genes per Cell | Reads Mapped to Genome (%) | Total Genes Detected | Median UMI Counts per Cell |
| --- | --- | --- | --- | --- | --- | --- |
| 1 | 1,460 | 623,754 | 3,529 | 96.0 | 26,874 | 13,890 |
| 2 | 3,797 | 309,348 | 2,892 | 96.7 | 27,044 | 9,946 |
| 3 | 2,520 | 656,319 | 2,243 | 95.6 | 26,402 | 7,067 |
| 4 | 3,336 | 171,518 | 3,147 | 95.8 | 27,066 | 10,216 |
| 5 | 618 | 162,440 | 3,714 | 96.0 | 22,387 | 16,654 |

Each cell was annotated to a cell type based on similarity in expression profiles with a reference dataset (Aran et al., 2019) (Figure S2B). Clusters were assigned a cell type based on the expression of previously reported markers for endometrial cell populations (Figure 2, C0-C16). Ten clusters were of epithelial origin (Clusters 2, 7, 13, 1, 5, 9, 14, 3, 15, 11) based on *EPCAM* expression (Figure S3A) and six of mesenchymal origin (Clusters 4, 6, 12, 0, 10, 8).

**Figure 2.**
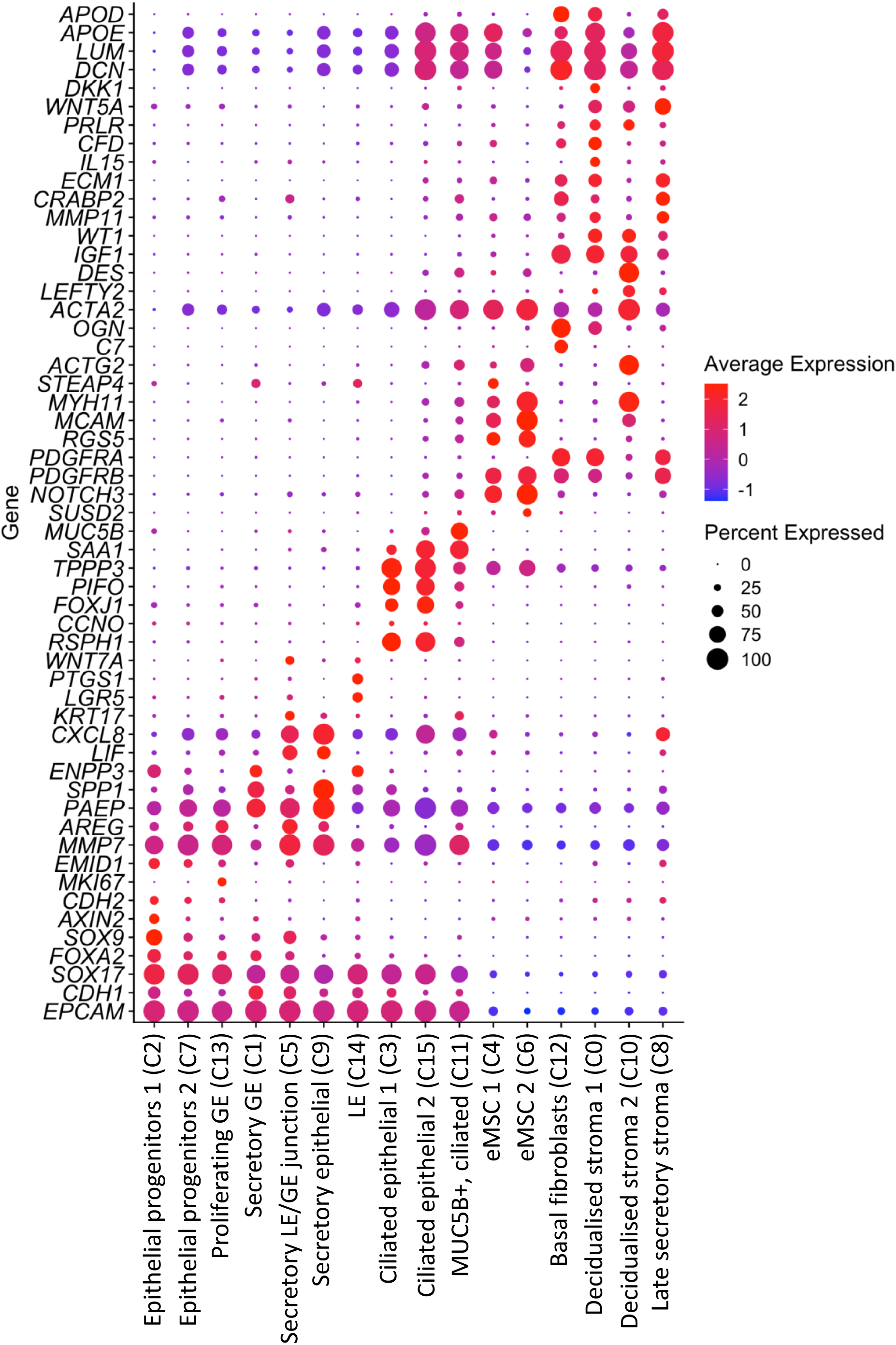
ScRNAseq identified 16 different endometrial cell populations. Dot plot showing previously published markers of different endometrial cell types used to annotate the cell populations.

Human endometrial stroma contains a subpopulation of perivascular cells fulfilling the classic MSC properties according to the International Society for Cell and Gene Therapy (Viswanathan et al., 2019), and by their stem cell functional properties (Masuda *et al*., 2012). Here we identified eMSCs on the basis of well characterised perivascular markers SUSD2 (Masuda *et al*., 2012; Sivasubramaniyan et al., 2013), MCAM (Crisan et al., 2008; Schwab and Gargett, 2007) and RGS5 (Bondjers et al., 2003). A total of 520 *SUSD2^+^* cells were identified (Table S2). *SUSD2* (Figure 1D blue dots, Figure 2), *MCAM* and *RGS5* (Figure 3A) expression was selectively enriched in cells in C6 and C4. A presumptive marker for endometrial perivascular cells from previous endometrial scRNAseq studies, *STEAP4* (Garcia-Alonso *et al*., 2021; Marečková *et al*., 2024), was confirmed in the C4 eMSC-1 but not C6 eMSC-2 subpopulations, but was also present in epithelial populations C1, C14 (Figures 2 and S3C). A previously assigned marker for myometrial perivascular cells, *MYH11* (Garcia-Alonso *et al*., 2021; Marečková *et al*., 2024) showed high expression in C6 eMSCs-1 but also in C10 decidualized stromal-2 cells (Figures 2 and 3). eMSCs were observed in all samples (Figure S3G, Tables S2 and S3).

**Figure 3.**
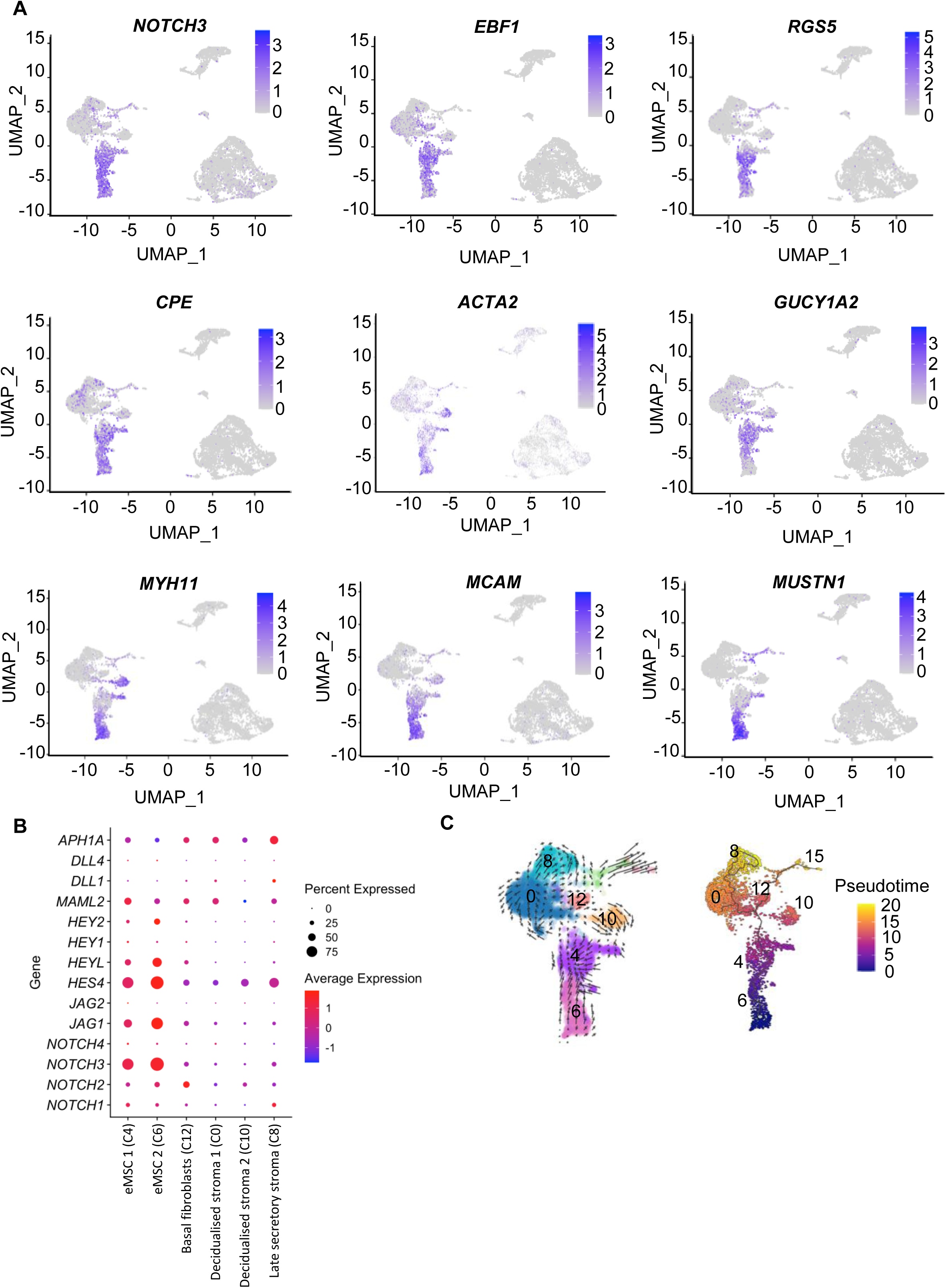
Two eMSC subpopulations in 6 mesenchymal cell populations identified in human endometrial stem/progenitor cell-enriched endometrial cells. **(A)** UMAP plots showing genes highly expressed in the 2 eMSC clusters (C4 and C6). **(B)** Dot plot showing NOTCH signalling genes in the endometrial mesenchymal cell populations. **(C)** Cell fate trajectory of mesenchymal populations; left panel, RNA velocity based on spliced and non-spliced transcripts using velocyto and scVelo. Velocity field projected onto UMAP overlayed with arrows showing the local average velocity evaluated on a regular grid. Right panel is the predicted cell fate trajectory using Monocle3 setting C6 as the root node containing the largest proportion of *SUSD2^+^MCAM*^+^ cells.

Typical mesenchymal genes*, PDGFRB* (Schwab and Gargett, 2007) and *ACTA2* were identified in all 6 mesenchymal clusters at varying expression levels (Figure 2). In contrast, expression of *PDGFRA* and *IGF1* was selectively enriched in the non eMSC clusters C12, C0, C10 and C8. We confirmed complement *C7* as a basal fibroblast gene (C12) (Figure 2) (Garcia-Alonso *et al*., 2021; Marečková *et al*., 2024). Decidualization, essential for embryo implantation, is a terminal differentiation of functionalis endometrial stromal cells into an epithelioid phenotype during the secretory stage of the menstrual cycle (Gellersen and Brosens, 2014). Two clusters of stromal cells expressing decidualization genes were identified, C0 and C10. Defining genes for C0 were *IL15* and the WNT inhibitor *DKK1,* and for C10 *LEFTY2* (Figure 2). Late secretory stroma (C8) is defined by high expression of *CXCL8*, but was also highly expressed in C9 and C5 secretory glandular and junctional epithelial cells, respectively (Figure 2). LEFTY2 is likely a negative regulator of the window of implantation and decidualisation and its high expression in C10 and C8 indicate that these stromal cell populations are of the later secretory phase (Salker et al., 2018).

Epithelial progenitors were then identified on the basis of their expression of previously identified specific protein markers that enriched for cells with functional stem/progenitor cell properties (Nguyen *et al*., 2017; Salamonsen *et al*., 2021). A total of 796 *EpCAM^+^CDH2^+^* epithelial progenitors were identified in 2 clusters (Table S4). Of these, 476 cells were *EpCAM^+^CDH2^+^SOX9^+^* (Table S5). *CDH2*, *SOX9* and *AXIN2* were concentrated in C2 and extended into C7 (Figures 1E, 2 and S3E). Clonogenic, self-renewing EPCAM^+^CDH2^+^SOX9^−^ cells were previously found in the basalis glands adjacent to the myometrium (Nguyen *et al*., 2017). By scRNAseq, rare populations of *CDH2^+^SOX9^−^* cells were scattered within several clusters (Figure 1E), mainly C2 and C7, but also in several stromal clusters and may represent *EPCAM^−^CDH2^+^* cells of stromal origin (Figure 1E). However, *EPCAM^+^CDH2^+^SOX9^+^* cells were almost exclusively localised to C2 and C7 (Figure 1E) suggesting this is a basalis glandular epithelial progenitor cell population which may express *CDH2* (N-cadherin), but not SOX9 protein. *EPCAM^+^CDH2^−^SOX9^+^* cells, representative of either luminal or glandular progenitor cells, were mostly present within C2 and C5. Mature epithelial *EPCAM*^+^*CDH2^−^SOX9^−^* cells were present within most epithelial clusters except C2. *EPCAM*^+^*CDH2^+^SOX9^+^* cells were mainly found in proliferative phase samples 1 and 2 (Supplemental Figure S3H).

*EPCAM^+^* populations with higher expression of the glandular epithelial marker, *FOXA2* (Kelleher et al., 2019) found in C2, C7, C13, C1 and C5 (Figures 2 and S3D) are likely of glandular epithelial origin. While C5 had higher expression of *FOXA2*, it also expressed secretory phase markers *LIF, PAEP* and *SPP1* and markers of luminal epithelium (*SOX9, KRT17, WNT7A, LGR5*) (Garcia-Alonso *et al*., 2021), suggesting a population in the junctional zone between the glandular (GE) and luminal epithelium (LE) (Figure 2). C9 and C14 had lower expression of *FOXA2*. C9 also expressed secretory markers suggesting another secretory LE subpopulation and C14, expressing *LGR5* and *ENPP3*, is potentially a mixture of secretory LE and LE cells (Figure 2). High *MKI67* expression in C13 indicates a proliferating GE population. C1 showed high expression of *FOXA2* and secretory and progesterone responsive markers *PAEP, SPP1* and *ENPP3* suggesting a secretory GE population (Figure 2). Epithelial C3, C15 and C11 with high proportions of cells expressing *EPCAM, FOXJ1* and *PIFO* were annotated as ciliated epithelial cells (Figure 2). C11 demonstrated high expression of *MUC5B* previously reported as defining an epithelial progenitor population (Tan et al 2022) but more likely endocervical cells (Marečková *et al*., 2024).

### eMSC populations identified in human endometrium

Highly enriched genes within eMSC C4 and C6 included classic MSC/eMSC signalling markers *NOTCH3* and *RGS5,* transcription factor*, EBF1,* and *CPE* which cleaves C-terminal basic residues of prohormones (Figure 3A). The *GUCY1A2* subunit involved in generating cGMP was expressed in 75% of C4 cells and just 11.6% of cells in all other clusters, highlighting its specificity for C4. MSC markers *MYH11 and MCAM* are present in C6 cells. *MUSTN1,* a development-dependent smooth and skeletal muscle regulatory gene involved in growth (Hadjiargyrou, 2018) showed the highest specificity of expression (93.9% vs 6.6%) and greatest fold change (3.16) in C6 cells (Figure 3A; Table S12). *NOTCH3* was most highly expressed in C6 *SUSD2*-expressing cells followed by C4 (Figure 3B, Table S10). *NOTCH3* (role in cell fate decisions) is most highly and almost exclusively expressed in most C4 and C6 eMSCs while *NOTCH1, 2,* and *4* are expressed more broadly (Figure 3B). Ligand *JAG1,* bHLH family transcription factor transcripts *HES4, HEYL*, *HEY2* and transcriptional co-activator *MAML2* and the γ-secretase subunit, *APH1A* are also expressed indicating active canonical NOTCH signalling (Weber et al., 2014) regulating cell fate decisions between eMSCs and various endometrial niche cells. Biological processes enriched for the genes in C4 eMSC include wound healing and vasculature development (Figure S4A). Genes expressed in C6 were also enriched for vasculature development, actin cytoskeletal organisation and muscle development. In contrast, the more mature decidualised stroma, C0 and late secretory stroma C8, were enriched for processes related to differentiated cells including cellular response to amino acid stimulus and extracellular matrix organisation (Figure S4B).

RNA velocity showed that eMSC C6 and C4 are the more immature cell types (Figure 3C), specifically highlighted for *SUSD2^+^* eMSCs (Figure S5A). These predicted progenitor cell populations, C4 and C6, were used as root populations to predict cell differentiation states using cell fate trajectory analysis showing that this cell state differentiated toward more mature fibroblasts in C0, C8 and C12 (Figure 3C). Cell cycle phase analysis revealed that most cells in each mesenchymal cell population were in the G1, with the least in C4 eMSC-1 followed by C6 eMSC-2 subpopulations suggesting eMSC are more proliferative than fibroblasts (Figure S5D).

### Endometrial epithelial progenitor populations identified in human endometrium

Key endometrial epithelial progenitor, GE and LE markers were examined in the epithelial subpopulations (Figure 4A). Highly enriched genes for the two epithelial progenitors, C2 and C7 included transcription factors *SOX9* and *SOX17* (Figures 2 and 4A), developmental genes *PRR15* and *CPM,* which cleaves basic C-terminal residues from peptides/proteins with potential roles in controlling peptide hormone and growth factor activity at the cell surface (Figure 4B). Genes with higher specificity to C2 than C7, include serine protease inhibitor *SERPINA5*, protease inhibitor *CST1,* thyrotropin releasing hormone and modulator of hair follicle mitochondrial growth *TRH*, and morphogen *IHH*. *TRH, CST1* and *IHH* were the most highly expressed genes in C2 (Table S8). High expression of steroid hormone receptors *PGR*, *ESR1* (Figure S7A) and their co-regulators (Figures S7B) in C2 and C7 was also notable.

**Figure 4.**
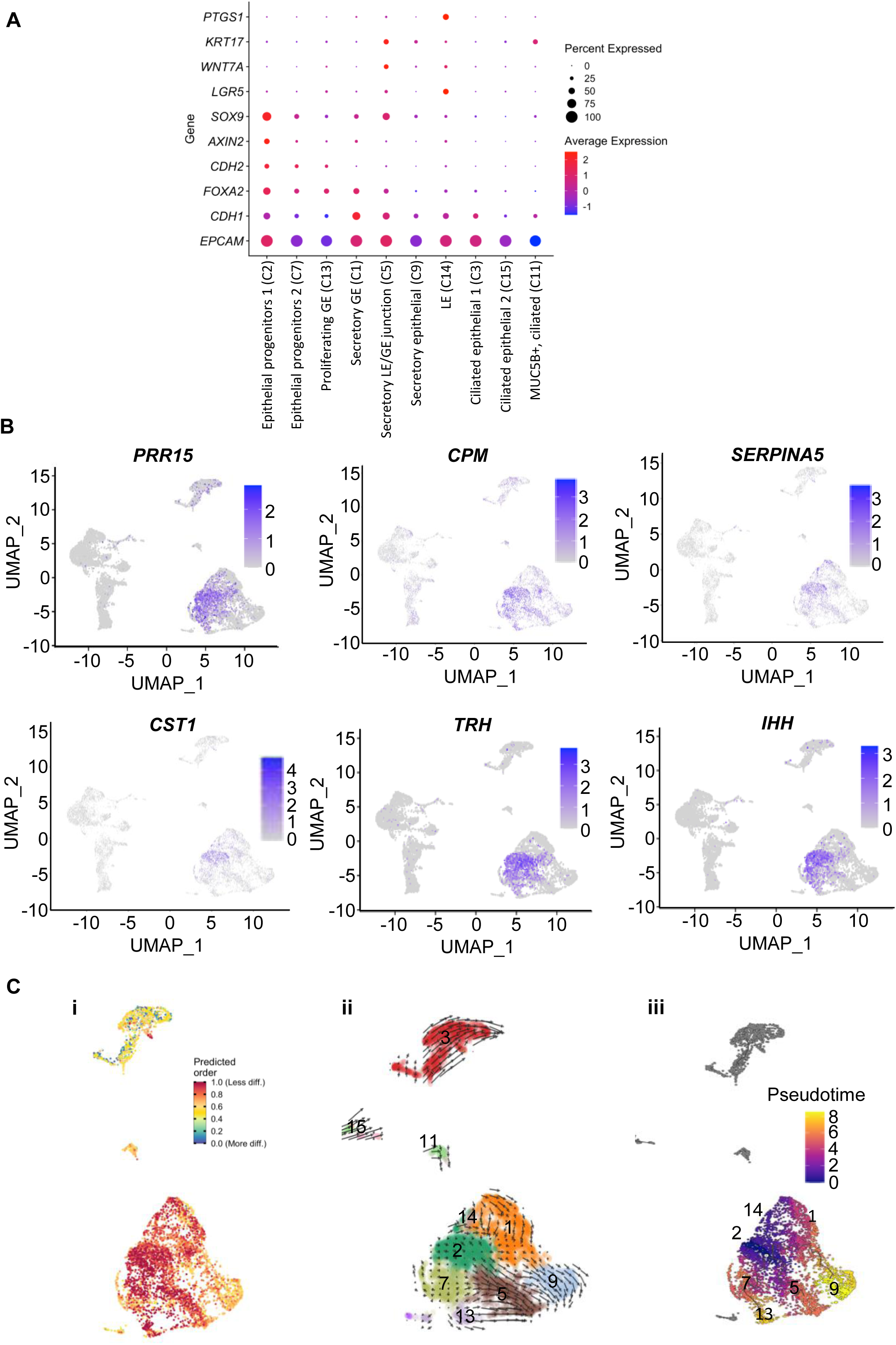
Two epithelial progenitor subpopulations in 10 epithelial populations identified in human endometrial stem/progenitor cell-enriched endometrial cells. **(A)** Dot plot showing the expression of endometrial epithelial and epithelial progenitor markers in the epithelial cell populations. C2 (Epithelial Progenitors-1) has high expression of *CDH2*, *SOX9*, *AXIN2* and *FOXA2*. **(B)** UMAP plots showing highly expressed genes in Epithelial Progenitor-1 and-2 (C 2 and C7 respectively). **(C)** Cell fate trajectory of epithelial cell populations. (i) UMAP depicting the distribution of CytoTRACE scores among epithelial cells. Green-blue indicates low stemness, more differentiated; red indicates high stemness, less differentiated. (ii) modelled RNA velocity based on spliced and non-spliced transcripts using velocyto and scVelo. Velocity field projected onto UMAP overlayed with arrows showing the local average velocity evaluated on a regular grid. (iii) predicted cell fate trajectory using Monocle3 and setting C2 as the root node containing the largest proportion of *CDH2^+^SOX9^+^* cells.

CytoTRACE predicted that C2 epithelial cells had the least differentiated state and transitioned to more mature epithelial populations, based on the number of detectable expressed genes/cell as a determinant of developmental potential (Figure 4Ci). RNA velocity showed that C2 and C7 are epithelial progenitor populations with more immature cell types (Figure 4Cii), specifically highlighted for *CDH2* and *SOX9* epithelial progenitors (Figure S5B, C). Cell fate trajectory analysis setting C2 and C7 as root populations predicted their differentiation toward more mature GE and LE cells in C1, C5, C9 and C13 (Figure 4Ciii). The majority of C2 epithelial progenitors-1 are in the G1 phase of the cell cycle with few in S phase (Figure S5D). The C7 epithelial progenitor-2 cells have more cells in the S phase, indicating increasing DNA replication, suggesting their role as transit amplifying cells with a more differentiated state than C2 cells within the epithelial cell hierarchy.

These data together with gene expression analysis suggested C2 as a basalis epithelial progenitor population. This was confirmed by TRH immunolocalisation in basalis GE and absence within the functionalis (Figure 5A) and similarly for IHH (Figure 6B). Members of the thyroid signalling pathway up- and down-stream of TRH were examined in all cell clusters (Figure 5B), showing high expression of *DIO2* in C2, C12, C0 and C8, *THRA* in C6 eMSC-2, C12 basal fibroblasts and C10 decidualised stroma and *NCOA1* also in those cell types. This suggests that TRH secreted from epithelial progenitors could be stimulating thyroid signalling in other cell types of the basal endometrium. Biological processes enriched for C2 genes included negative regulation of apoptotic signalling and gland development and in C7 for signal transduction, gland development and regulation of cell proliferation (Fig S6).

**Figure 5.**
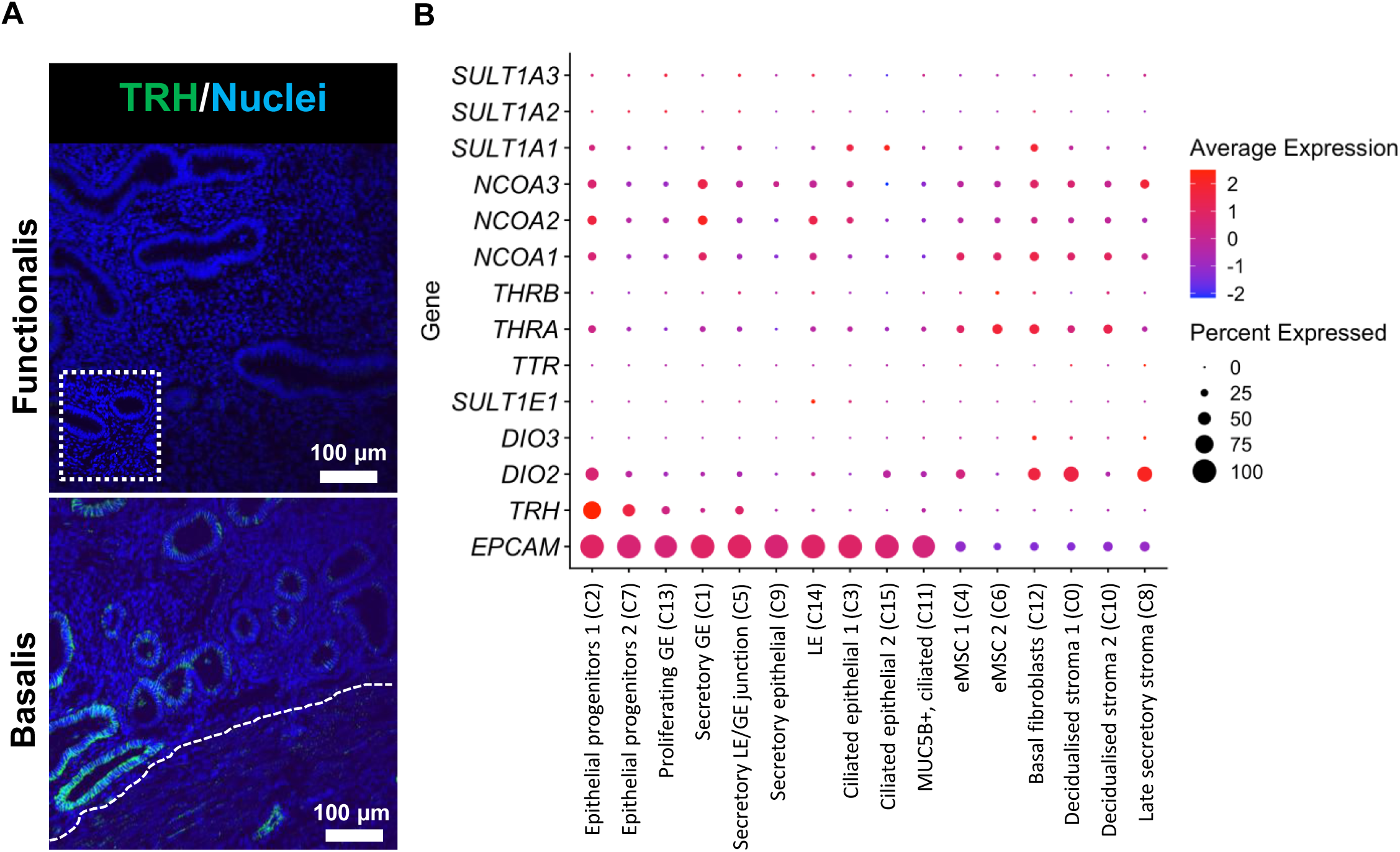
TRH protein and gene expression in human endometrium. **(A)** Immunolocalization of TRH to basalis glands and absence in the functionalis in full thickness endometrium. Representative image of n=5 proliferative and n=3 secretory phase endometrial tissues. White dashed line, endometrial-myometrial border; white dotted line, isotype control inset. Scale bar, 100 µm. **(B)** Dot plot showing genes involved in thyroid signalling and their expression in the 16 endometrial cell populations.

**Figure 6.**
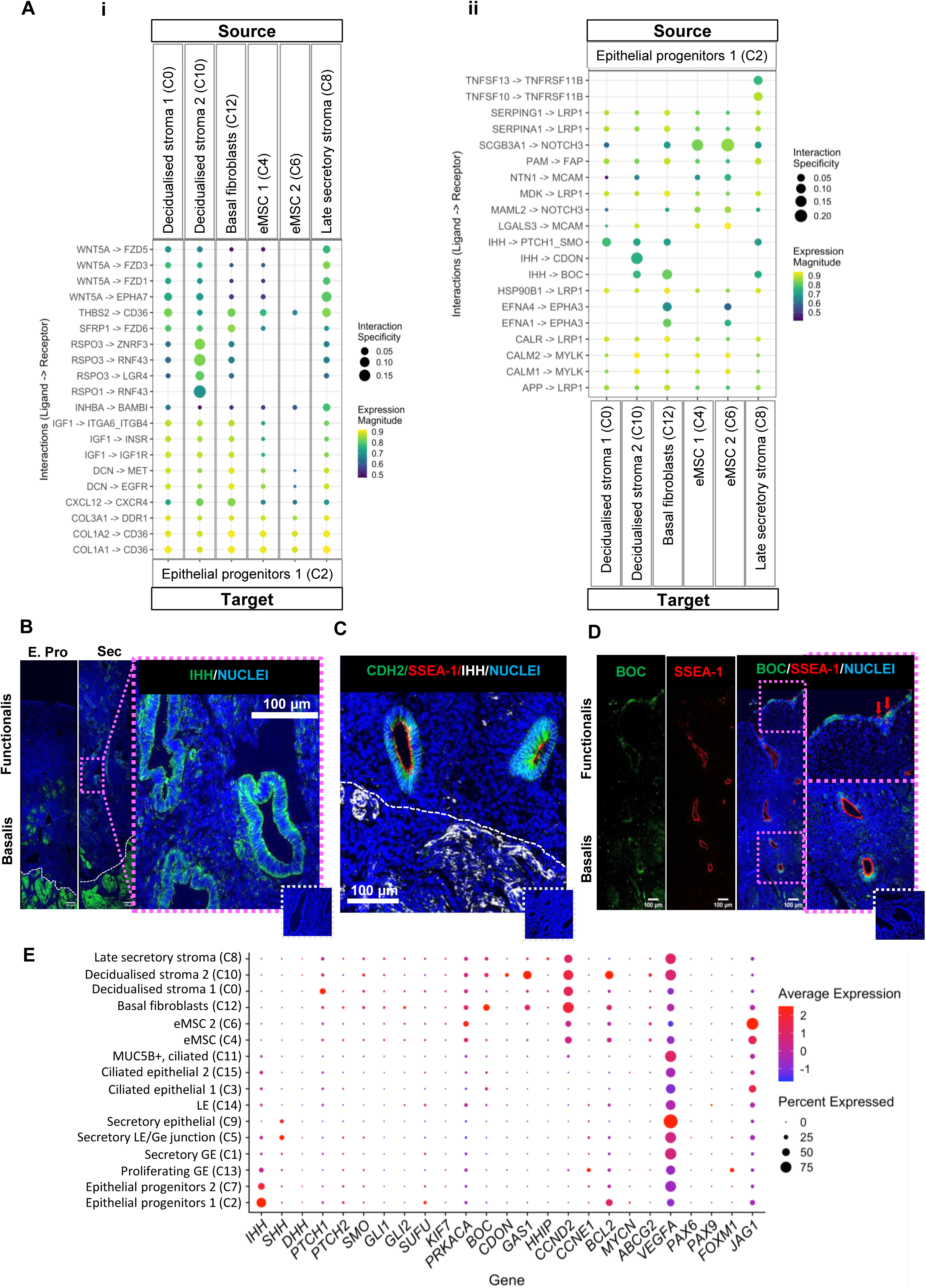
Ligand-receptor interactions between human endometrial epithelial progenitor cells and mesenchymal cells. **(A)** The top 20 interactions between ligands based on rank aggregate from (i) mesenchymal populations and receptors found on epithelial progenitors (ii) epithelial progenitors (C2) and receptors found on mesenchymal cell populations. The expression magnitude as estimated by SingleCellExperiment’s LRScore and interaction specificity as determined using NATMI’s edge specificity weights from 0 to 1, where 1 means both the ligand and receptor are uniquely expressed in a given pair of cells. **(B)** Immuno-localization of IHH in proliferative (Pro) and secretory (Sec) phase endometrium. **(C)** Immuno-colocalization of CDH2, SSEA1 and IHH in basalis glands. **(D)** Immuno-colocalization of BOC and SSEA1 in full thickness endometrium in luminal epithelium (upper inset) and basalis glands (lower inset). Red arrows show SSEA1 LE immunostaining (B)-(D) Representative examples of n=5 proliferative and n=3 secretory phase endometrial tissues. Scale bar, 100 µm. White dotted line, isotype control inset; white dashed line, endometrial-myometrial border. **(E)** Dot plot showing expression of genes involved in hedgehog (HH) signalling in endometrial cell populations.

### Endometrial epithelial stem/progenitor cell interactions in the basalis niche

To determine the cell-cell interactions between the different cell populations of the endometrium and to define the endometrial epithelial progenitor cell niche, LIANA was used, revealing the potential role of the WNT signalling pathway in maintaining this niche. Strong signalling interactions were predicted between the late secretory stromal and decidualized stromal cell populations expressing *WNT5A* and *SFRP1* (C8, C0, C10) and basalis C2 epithelial progenitors-1 expressing *FZD5, 3, 1* and *6* (Figure 6Ai). Similarly, WNT signalling may be potentiated through interactions between C10, C8 and C12 niche cells expressing *RSPO-1* and -*3* and C2 epithelial progenitors-1 expressing *RNF43, ZNRF3* and *LGR4*. Other predicted interactions were identified between IHH ligand from C2 epithelial progenitors-1 and receptors PTCH1/SMO on C12 basal stromal fibroblasts, C0 decidualised stroma-1, C10 decidualised stroma2, and C8 late secretory stroma (Figure 6Aii). Similarly, *IHH* expressed in C2 epithelial progenitors-1 interacts with Hedgehog (HH) co-receptors *CDON* and *BOC* expressed in C10 decidualised stroma-2 population, and *BOC* in C12 basalis stromal fibroblasts and C8 late secretory stroma (Figure 6Aii). IHH immunolocalized to endometrial basalis epithelium, some stromal cells and myometrium of the proliferative phase uterus, and in the basalis and functionalis epithelium, stroma, and myometrium of the secretory phase (Figure 6B). The epithelial progenitor markers CDH2 and SSEA1 co-localised with IHH in basalis GE (Figure 6C). In the human endometrium, BOC and SSEA1 co-localised in the basalis GE and LE corresponding with the potential for these proteins having a role in cell migration signalling (Kim et al., 2020) (Figure 6D).

Analysis of HH signalling in the identified endometrial cell populations highlighted that *IHH* is the main HH member in the endometrium and most expressed in epithelial progenitors (Figure 6E). *SHH* is only weakly expressed in secretory epithelial cells (Figure 6E). In addition to *BOC*, *CDON* and *PTCH1*, other genes involved in HH signalling found in endometrial cell populations include *JAG1*, *VEGFA*, *BCL2*, *CCND2*, *GAS1* and *GLI1 AND 2*, mostly in stromal populations, suggesting potential signalling interactions between epithelial progenitors and stromal cells (Figure 6E).

## Discussion

This study showed that the human endometrium contains at least four major stem/progenitor cell states of epithelial and mesenchymal origins that transition to mature cell states to regenerate and repopulate the endometrial functionalis following menstruation. Most previous scRNA seq analyses of the human endometrium have been unable to identify human endometrial epithelial progenitor cell populations because they have not sampled basalis and/or enriched cell populations using known stem/progenitor markers (Fonseca *et al*., 2023; Garcia-Alonso *et al*., 2021; Lv *et al*., 2022; Tan *et al*., 2022; Wang *et al*., 2020). Here, we showed the efficacy of our novel approach for characterising the rare human endometrial stem/progenitor populations and their niches by first enriching them by FACSorting using published markers of functional endometrial stem/progenitor cells and performing scRNA seq analysis. We identified 796 *EpCAM^+^CDH2^+^* epithelial progenitors separating into 2 clusters from 11,731 cells from the five patient samples. Of these, 476 also expressed *SOX9* (*EpCAM^+^CDH2^+^SOX9^+^*), mainly in the same 2 clusters. In comparison, the Human Endometrial Cell Atlas (HECA) (Marečková *et al*., 2024) comprising >313,000 cells identified 359 *CDH2^+^SOX9^+^* basalis epithelial cells from 8 of 63 patients, only 3 cells total from 2 hysterectomies and 342 cells from a single biopsy. In addition to these 4 stem/progenitor populations, we also identified 12 other populations of endometrial cell types.

Our single cell analysis confirmed two major populations of ciliated epithelial cells distinct from other cell types (Garcia-Alonso *et al*., 2021; Wang *et al*., 2020) however, the two populations in our study had low expression of the GE marker, *FOXA2*. We agree with others (Marečková *et al*., 2024) that the previously annotated *MUC5B^+^* population as a progenitor population (Tan *et al*., 2022) is unlikely given the absence of *CDH2* expression as identified in our study. Rather these ciliated *MUC5B^+^* cells are more likely endocervical cells given their high expression of genes/proteins also found in the cervix as predicted previously (Marečková *et al*., 2024). We identified four distinct GE populations assigned according to their *FOXA2* expression, including a proliferating population with high *MKI67* expression, secretory GE and two epithelial progenitor populations. For the first time, we identified a population indicative of a group of secretory cells found in the junctional zone between the GE and LE (at gland opening) due to their high expression of *FOXA2* and LE markers *KRT17* and *WNT7A* and secretory markers *CXCL8*, *PAEP* and *LIF*. One secretory epithelial population with no clear luminal or epithelial markers but high expression of *LIF*, *PAEP* and *SPP1, WNT7A* and *LGR5* indicating a LE subpopulation, which may be potential luminal progenitors due to *SOX9^+^* expression (Garcia-Alonso *et al*., 2021; Tan *et al*., 2022). The LE is likely derived from SOX9^+^ basalis epithelial cells (Salamonsen *et al*., 2021).

Our study identified two distinct perivascular eMSC subpopulations expressing classic MSC markers, including SUSD2. Both express *NOTCH3* and signalling pathway members as previously reported (Garcia-Alonso *et al*., 2021; Marečková *et al*., 2024) as do cultured human SUSD2^+^ eMSC that undergo decidualization differentiation via this pathway (Murakami et al., 2014). *MUSTN1* was highly selectively enriched for eMSC-2 (C6) and *GUCY1A2* for eMSC-1 (C4), signature markers not previously described for eMSC. eMSC-2 are likely the more immature, perivascular cells expressing similar genes (*MYH11, SOD2* and *SLIT3*) as the Pericyte-2 population (Queckbörner *et al*., 2021), and myometrial SMC (*ACTA2*) and mPV clusters of the HECA (Marečková *et al*., 2024). We suggest that the mSMC and mPV cells in the HECA are more likely eMSC than myometrial cells as the myometrium is not sampled in biopsies and poorly dissociates using endometrial processing protocols for hysterectomies. eMSC-1 (C4) have a closer profile to Pericyte-1 (Queckbörner *et al*., 2021) and both ePV1b and ePV2 of the HECA. Despite varying terminologies used, considerable consensus on eMSC gene profiles exists between published endometrial studies.

We identified two major endometrial epithelial progenitor cell states. They expressed key genes previously identified as protein markers of endometrial epithelial progenitor cells; *CDH2*, *AXIN2* and *SOX9*. Our methods allowed us to fully characterise these two populations to generate a human endometrial epithelial progenitor molecular signature. Recently, *CDH2^+^* epithelial cells were identified in a scRNA seq analysis using a much greater number of samples (n=63) including three hysterectomy samples (Marečková *et al*., 2024), highlighting the difficulty in capturing progenitor cells from biopsies but also the strength of our methodology employed when just five samples were used. Both endometrial epithelial progenitor populations showed high *ESR1* and *PGR* expression correlating with their protein localisation in basalis endometrium throughout the menstrual cycle. In contrast, functionalis ESR1 and PGR localisation in GE and LE vary across the menstrual cycle in response to circulating estrogen and progesterone (Gargett et al., 2008). While functionalis epithelium proliferates and differentiates respectively in response to these hormones, no such response occurs in the basalis. Rather, non-proliferating SSEA1^+^SOX9^+^ epithelial cells rapidly migrate from the protruding gland stumps on the first 2-3 days of menses (Salamonsen *et al*., 2021).Once the endometrium has resurfaced, epithelial cells in the neck of the glands proliferate independently of estrogen on day 3-4 of menses, but, the N-cadherin^+^ deep basal region of those same glands does not despite high ESR1 expression (Ferenczy et al., 1979). We identified expression of the nuclear coactivators (*NCOA2, 7*) and corepressors (*NCOR1, 2*) that activate and inhibit ESR1 transcription, respectively in the epithelial progenitor clusters of basalis epithelium. These corepressors may be responsible for the very low proliferative response in SSEA1^+^SOX9^+^ basalis epithelial and stromal cells. The role of coactivator/repressor interactions in the dichotomous response between functionalis and basalis endometrial cell ESR1 is currently unknown, but our findings suggest future research avenues.

*TRH* expression in the endometrium has been little studied and yet its high specificity in basalis epithelial progenitors suggests a role for thyroid hormone signalling in endometrial development and monthly tissue regeneration. TRH acts on anterior pituitary TRHR to release TSH which stimulates T3 and T4 release from the thyroid gland (Brown et al., 2023) but its role in human endometrium remains unknown. TRH has been identified in the proliferative phase and in postmenopausal endometrium (Zieba et al., 2015). Our earlier gene profiling studies showed that post-menopausal endometrium has a similar gene expression profile as pre-menopausal basalis (Nguyen *et al*., 2012). The macaque uterus expresses all the receptors involved in thyroid hormone function suggesting that the human uterus is likely similar with local production and regulation of thyroid signalling (Hulchiy et al., 2012). Patients with endometriosis and infertility have increased serum prolactin levels following TRH treatment (Cunha-Filho et al., 2002). Indeed, infertile women with hypothyroidism have higher PRL levels associated with abnormal menstrual cycle patterns (Fröhlich and Wahl, 2019). The expression of *TRH* and its receptors in the same tissue suggests local paracrine action, although *TRHR* was not detected in our data or the HECA. However, we found *DIO2*, which converts the prohormone T4 to active T3 (Ambrosio et al., 2017), expression in the epithelial progenitors, basal fibroblasts, decidualized stroma and late secretory stromal cells suggesting T4 to T3 conversion maintains endometrial homeostasis. DIO2 is highly expressed in differentiating myoblasts and has a role in differentiating activated satellite (stem) cells.

Our two endometrial epithelial progenitor populations had enriched expression of *IHH*, also confirmed at the protein level. Hedgehog signalling is a key developmental pathway in vertebrates (Sigafoos et al., 2021) and IHH is 1 of 3 hedgehog ligands that binds patched receptors (PTCH1, PTCH2) near the primary cilium present on most cells, including human endometrial stromal cells. Here it alleviates PTCH inhibition of SMO to activate downstream transcription factors GLI1 (Walterhouse et al., 2003). In mouse uteri, Ihh regulates progesterone signalling between the epithelium and stroma (Lee et al., 2006). Conditional Smo knockout leads to shorter diestrus, uteri not thickening in estrogen dependent cycle stages, abnormal endometrial morphology and infertility (Roberson et al., 2023). Downregulated IHH has been linked to human adenomyosis (Zhou et al., 2021), where endometrial basalis grows into myometrium. We identified cell interactions between epithelial progenitor IHH and basal stromal IHH co-receptors CDON and BOC, PTCH1, and downstream signalling molecules including the GLIs. In mice, primordial germ cells migrating from the hindgut to the genital ridges express both SSEA1 and Boc (Kim *et al*., 2020), suggesting a potential role for the IHH co-receptors in migration and re-epithelisation of epithelial progenitors over the raw endometrium as it repairs during menses (Salamonsen *et al*., 2021). Together, this suggests a role for IHH in regulating autocrine/paracrine endometrial epithelial migration and repair during menses, regeneration, and potential maintenance of the stem cell niche.

Limitations that may impact interpretation and generalisability of our findings include the relatively small sample size and inherent variability among patient samples, including differences in menstrual cycle stage, age, and pathologies, which could influence cell composition and gene expression profiles. The choice of clustering resolution, while computationally optimised and biologically informed, may not fully align with all biologically meaningful cell types and states. These observations highlight the need for validation in larger, more diverse datasets to confirm the robustness and broader applicability of the results.

Characterising gene expression profiles of endometrial stem/progenitor cell types and states at the single cell level and further examining their niche cells will elucidate their role in regenerating endometrium each menstrual cycle and in the pathogenesis of endometrial proliferative disorders such as endometriosis, adenomyosis, Asherman’s disease and thin endometrium, possibly identifying new targets for developing treatments for these intractable diseases.

## Experimental Procedures

### Sample collection

Full thickness endometrial tissues containing the functionalis and basalis were collected from women <50 years undergoing hysterectomy (n=5) for benign uterine conditions including adenomyosis, endometriosis and fibroids (Table 1). Exclusion criteria were previous endometrial ablations, thickened endometrium, endometrial cancer and taking hormone therapy. Eligible women gave written informed consent. Human ethics approval was obtained from Monash Health HREC and the US Department of Defence OHRO (RES-19-0000-388A/E00700.1a.1b). Endometrial samples were either fixed in 4% PFA overnight at 4C (n=3) or placed into cold DMEM:F12 collection medium supplemented with 5% newborn calf serum and antibiotics/antimycotic and immediately transported at 4C to the laboratory for processing (n=5).

### Isolation of primary endometrial cells

Endometrial tissue was digested to single cells within 18 hours. The functionalis was removed by scraping the tissue off the surface of the myometrium and finely minced, then the full basalis was recovered by dissecting 1-2 mm myometrium with attached endometrium (visible microscopically) and finely dicing the tissue (Nguyen *et al*., 2017) (Figure 1A).The minced tissue underwent a 2-step enzymatic/mechanical dissociation with vigorous trituration every 5 min at 37C in PBS (Gibco) containing collagenase type I (5 mg/mL; Worthington), DNAse I (0.08 mg/mL; Worthington) and glucose (5mM; Merck), then strained (40 µM sieve, Falcon) to separate stromal cells from epithelial gland fragments. Red blood cells in the stromal fraction were removed by Ficoll-paque (Sigma-Aldrich) density gradient separation. Epithelial cells underwent a second digestion with a similar cocktail using collagenase type II (0.8 mg/ml, Worthington) and DNAse I 0.16 mg/mL, then epithelial cells were strained using a 40 µM sieve. The single cell fractions were counted and combined for FACS staining.

### 6-way FACSorting to enrich endometrial cells with stem/progenitor cell subpopulations

Single cell suspensions were blocked with Fc Receptor blocker (FcR; 2 µL/10^6^ cells, Miltenyi Biotec) and Rat IgG (4.4 µg/10^6^ cells; Jackson ImmunoResearch) as described (Wyatt et al., 2021). Cells were then distributed in tubes as unstained, viability and Fluorescence minus 1 (FMO) controls (up to 5X10^5^ cells in 20 µL) and the full antibody panel (remaining cells at 5X10^5^/20 µL at the same final antibody concentration) and incubated with conjugated antibodies, PE-CD325 (N-cadherin) (10 µg/mL), BV421-CD15 (SSEA1) (1.25 µg/mL; BioLegend), APC-SUSD2 (1.25 µg/mL; BioLegend,), BB515-CD326 (EpCAM) (0.25 µL/5X10^5^ cells, BD Biosciences), PECy7-CD31 (PECAM1, endothelial exclusion marker) (0.25 µl/5X10^5^ cells, BD Biosciences) and CD45 (pan leukocyte exclusion marker) PECy7-CD45 (0.25 µL/5X10^5^cells, BD Biosciences), for 25 min at 4°C. Propidium iodide (PI; 62.5 ng/mL; BD Biosciences) was added 10 min before sorting (Wyatt *et al*., 2021). Compensation Plus Particle Set beads (60 µL/test; BD Biosciences) were stained as single staining protocols with the above antibodies following manufacturer’s instructions. Labelled endometrial cells underwent FACSorting of endometrial stem/progenitor cell subpopulations and their differentiated progeny (epithelial (EpCAM+) progenitors N-cadherin^+^/SSEA1^−^; N-cadherin^+^/SSEA1^+^; N-cadherin^−^/SSEA1^+^; mature epithelial cells, EpCAM^+^/N-cadherin^−^/SSEA1^−^) (Nguyen *et al*., 2017) and eMSC (SUSD2^+^; SUSD2^−^ fibroblasts) (Masuda *et al*., 2012) using an Influx Cytopeia-BD 5 laser sorter and BD FACS™ Sortware software (Monash University FlowCore). The gating strategy for sorting 6 cell subpopulations from viable PI^−^CD45^−^ cells is shown in Figure 1A and Fig S1.

### Single-cell RNA-sequencing

The 6 cell fractions were tagged using TotalSeq A anti-human hashtags. The stem/progenitor cell populations were pooled and mature epithelial and stromal cells added for scRNAseq for each of the 5 biological samples (Fig 1A) to generate barcoded single-cell 3′ gene expression libraries with the Chromium Single-cell 3′ GEM, Library & Gel Bead Kit v3 (10x Genomics) and their matched HTO library to resolve the hashtags (BioLegend). The quality of the libraries was assessed by Agilent Bioanalyzer High Sensitivity DNA chip. Libraries were sequenced to an average depth of 384,676 reads/cell on the Illumina NovaSeq6000 at UQ. Sequence quality checks revealed that HTO library construction had failed; and were excluded from subsequent analyses. Cell type identification was instead performed using the bioinformatic approaches described below.

### Processing scRNAseq data

Raw sequencing data was processed to fastq files using the mkfastq function in Cell Ranger (v6.0.1). Reads were aligned to the hg38 human reference, filtered for valid cell barcodes and unique molecular identifiers (UMIs) and counted using the count function. Doublets were identified in individual samples using the doubletFinder package in R (v4.1.2) (McGinnis et al., 2019) and removed. Single cells were then analysed using Seurat (v4.1.1) (Hao et al., 2021) to remove cells with very low numbers of expressed genes (<200), >25% mitochondrial genes and genes expressed in <3 cells. scTransform (Hafemeister and Satija, 2019) normalization (Choudhary and Satija, 2022) and Harmony (Korsunsky et al., 2019) was used to correct between sample variation due to technical and biological differences.

Normalised and batch corrected data were used as input for linear dimensional reduction and Principal Component Analysis (PCA). Cells were clustered using the Louvain algorithm and the first 20 principal components in Seurat. Clustering was performed across multiple resolutions between 0.01 and 1.0 and cluster stability was assessed using clustree (v0.5.1).

### Statistics

Differentially expressed genes (DEGs) between clusters were identified using the ‘FindMarkers’ function in Seurat using a non-parametric Wilcoxon rank sum test and restricting tests to genes expressed in at least 25% of cells in either of the two populations being compared. Expression in each cluster was compared to all other clusters combined. Gene expression differences were considered significant if they had an absolute log-fold change of ≥0.1 and adjusted p-value <0.05.

### Trajectory and RNA velocity analysis

Transcriptional profiles were used to estimate the transition between cell states in the endometrium using cell fate trajectory analysis in Monocle3 (Cao et al., 2019). Root cells were set computationally as the trajectory graph node containing the highest proportion of cells from the ‘earliest time point’. For cells of mesenchymal origin this was selected as the node where the most SUSD2^+^ cells were expressed. Similarly, the ‘earliest time point’ for cells of epithelial origin was selected as the node with *CDH2^+^SOX9*^+^, *CDH2^−^SOX*^+^ or *CDH2^+^SOX^−^EPCAM*^+^ as the root cells. RNA velocity was also used to assess cell trajectory using dynamic modelling with velocyto (La Manno et al., 2018) and scVelo (Bergen et al., 2020). Cellular Trajectory Reconstruction Analysis using gene Counts and Expression (CytoTRACE) predicted the differentiation state of the cells based on the number of detectably expressed genes per cell as a determinant of developmental potential. Raw single-cell expression counts were used as input in the ‘CytoTRACE’ function in the CytoTRACE R package v0.3.3 (Gulati et al., 2020) and results visualised using the ‘plotCytoTRACE’ function and coordinates generated from UMAP.

### Cell Interaction analysis

To investigate interactions between epithelial progenitors and mesenchymal cell types we performed cell communication analyses using LIANA (LIgand-receptor ANalysis frAmework) (Dimitrov et al., 2022). LIANA combines multiple tools and resources and provides a rank aggregate from the results of different methods serving as a consensus across methods. Analyses were run using epithelial progenitors, C2, as the target and mesenchymal clusters as the source and again with epithelial progenitors as the source. The top 20 interactions based on the rank aggregate were plotted.

### Experimental Procedures

The procedures describing cell type annotation, gene ontology analysis and immunofluorescence are described in the supplemental experimental procedures.

## Resource Availability

### Lead contact

Further information or requests should be directed to and will be fulfilled or facilitated by the Lead Contact, Caroline Gargett

### Materials availability

This study did not generate new unique reagents.

### Data code and availability

All scRNAseq data generated in the present study will be deposited at the NCBI Gene Expression Omnibus and will be publicly available at the date of publication. No original code and/or algorithms are reported: however, code used for the data analysis can be provided on request. Any additional information required to reanalyse the data reported in this paper is available from the lead contact on request.

## Acknowledgements

This work was supported by US Department of Defense Award No. W81XWH1910364/PR180827 (CEG, GWM, LR) and National Health and Medical Research Council (Australia) Investigator Fellowship Nos 1173882 (CEG) and 1177194 (GWM). The authors acknowledge research nurses Jenny Ryan and Madi Bates, clinician Dr Kate Tyson and women who donated tissue. The authors acknowledge use of the services and facilities of Micromon Genomics at Monash University, Monash University Flowcore and Monash Histology Platform at Hudson Institute of Medical Research. We thank Anjali Henders, Leanne Wallace, and the staff of the Human Studies Unit at the Institute for Molecular Bioscience for support with recruitment, sample processing and genotyping.

## Author contributions

CEG and GWM conceptualised and together with HCF, SM and CEF designed the research. HCF, SM, CEF, KAW, FLC and EMG performed the research; HCF and SM analyzed the data; HCF, SM and FLC visualized the data and results; HCF, SM, BM, CEG and GWM interpreted results; CEG was the project administrator; CEG, JT, GWM contributed resources; HCF, SM and CEG wrote the paper; All authors reviewed and edited the paper.

