## Supplementary Information for "Molecular signature of human endometrial stem/progenitor cells at the single cell level"

**Supplementary Material**

#### Supplemental Tables

**Table S1**. Patient characteristics of the five samples subjected to scRNA-seq.

| Donor ID | Age | Cycle phase | Pathologies |
|---|---|---|---|
| 1 | 45 | Proliferative | Fibroids |
| 2 | 41 | Proliferative | Adenomyosis |
| 3 | 29 | Periovulatory | Endometriosis, adenomyosis |
| 4 | 48 | Secretory | Adenomyosis, fibroid |
| 5 | 46 | Secretory | Fibroids |

Cycle phase was determined from progesterone (P4) levels, proliferative (0.18-2.84 nM/L), periovulatory (2.84-5.82 nM/L), secretory (5.82-75.9 nM/L).

**Table S2.** The number of *SUSD2*$^+$ cells in each sample analysed by scRNA seq.

| Donor ID | Cycle phase | ***SUSD2***$^+$ |
|---|---|---|
| 1 | Proliferative | 157 |
| 2 | Proliferative | 145 |
| 3 | Periovulatory | 47 |
| 4 | Secretory | 132 |
| 5 | Secretory | 39 |

**Table S3.** The number of *SUSD2*$^{+}$ cells in each cluster analysed by scRNA seq.

| **Cluster** | ***SUSD2***$^{+}$ |
|---|---|
| 0 | 24 |
| 1 | 23 |
| 2 | 22 |
| 3 | 9 |
| 4 | 117 |
| 5 | 15 |
| 6 | 220 |
| 7 | 2 |
| 8 | 24 |
| 9 | 0 |
| 10 | 0 |
| 11 | 37 |
| 12 | 16 |
| 13 | 3 |
| 14 | 4 |
| 15 | 4 |

**Table S4.** The number of *EPCAM*$^{+}$*CDH2*$^{+}$ cells in each sample analysed by scRNA seq.

| **Donor ID** | **Cycle phase** | ***EPCAM***$^{+}$***CDH2***$^{+}$ |
|---|---|---|
| 1 | Proliferative | 26 |
| 2 | Proliferative | 625 |
| 3 | Periovulatory | 67 |
| 4 | Secretory | 76 |
| 5 | Secretory | 2 |

**Table S5.** The number of *EPCAM*$^{+}$*CDH2*$^{+}$*SOX9*$^{+}$ cells in each cluster analysed by scRNA seq.

| Cluster | *EPCAM*$^{+}$*CDH2*$^{+}$*SOX9+* |
|---|---|
| 0 | 0 |
| 1 | 28 |
| 2 | 279 |
| 3 | 2 |
| 4 | 0 |
| 5 | 48 |
| 6 | 0 |
| 7 | 95 |
| 8 | 1 |
| 9 | 1 |
| 10 | 0 |
| 11 | 0 |
| 12 | 0 |
| 13 | 17 |
| 14 | 5 |
| 15 | 0 |

**Table S6.** Top 30 genes in cluster 0

| Gene | Protein name | Fold Change | pct.1 | pct.2 | p_val_adj |
|---|---|---|---|---|---|
| IGF1 | Insulin Like Growth Factor 1 | 2.41 | 0.828 | 0.134 | 0 |
| PTGDS | Prostaglandin D2 Synthase | 2.27 | 0.757 | 0.358 | 0 |
| CFD | Complement Factor D | 2.23 | 0.565 | 0.097 | 0 |
| DCN | Decorin | 2.21 | 0.997 | 0.475 | 0 |
| SERPINF1 | Pigment epithelium-derived factor | 2.12 | 0.921 | 0.321 | 0 |
| H19 | | 2.11 | 0.772 | 0.086 | 0 |
| MEG3 | | 2.10 | 0.848 | 0.142 | 0 |
| LUM | Lumican | 2.07 | 0.986 | 0.481 | 0 |
| PDGFRA | Platelet Derived Growth Factor Receptor Alpha | 2.05 | 0.821 | 0.102 | 0 |
| COL3A1 | Collagen Type III Alpha 1 Chain | 2.03 | 0.997 | 0.574 | 0 |
| SFRP4 | Secreted Frizzled Related Protein 4 | 2.01 | 0.737 | 0.131 | 0 |
| RORB | RAR Related Orphan Receptor B | 2.01 | 0.782 | 0.098 | 0 |
| RAMP1 | Receptor Activity Modifying Protein 1 | 1.94 | 0.937 | 0.399 | 0 |
| COL1A1 | Collagen Type I Alpha 1 Chain | 1.90 | 0.997 | 0.578 | 0 |
| APOD | Apolipoprotein D | 1.90 | 0.623 | 0.077 | 0 |
| IGFBP2 | Insulin Like Growth Factor Binding Protein 2 | 1.87 | 0.966 | 0.615 | 0 |
| C1R | Complement C1r | 1.86 | 0.933 | 0.487 | 0 |
| PCOLCE | Procollagen C-Endopeptidase Enhancer | 1.86 | 0.904 | 0.476 | 0 |
| VCAN | Versican core protein | 1.85 | 0.887 | 0.529 | 0 |
| PTN | Pleiotrophin | 1.85 | 0.787 | 0.274 | 0 |
| APOE | Apolipoprotein E | 1.84 | 0.908 | 0.454 | 0 |
| APCDD1 | APC Down-Regulated 1 | 1.83 | 0.61 | 0.129 | 0 |
| NR2F1 | Nuclear Receptor Subfamily 2 Group F Member 1 | 1.83 | 0.66 | 0.143 | 0 |
| C1S | Complement C1s | 1.80 | 0.871 | 0.346 | 0 |
| HAND2-AS1 | | 1.79 | 0.66 | 0.099 | 0 |
| ISLR | Immunoglobulin superfamily containing leucine-rich repeat protein | 1.77 | 0.801 | 0.182 | 0 |
| MME | Membrane metalloendopeptidase | 1.75 | 0.608 | 0.057 | 0 |
| HAND2 | Heart and neural crest derivatives expressed 2 | 1.75 | 0.582 | 0.097 | 0 |
| HELLPAR | | 1.75 | 0.626 | 0.088 | 0 |
| CPXM1 | Probable carboxypeptidase X1 | 1.71 | 0.761 | 0.133 | 0 |

**Table S7.** Top 30 genes in cluster 1.

| Gene | Protein name | Fold change | pct.1 | pct.2 | p_val_adj |
|---|---|---|---|---|---|
| MT1G | Metallothionein 1G | 2.37 | 0.818 | 0.395 | 0 |
| SCGB1D2 | Secretoglobin family 1D member 2 | 2.22 | 0.766 | 0.512 | 3.33E-183 |
| LINC01320 | | 2.02 | 0.934 | 0.452 | 0 |
| MT1F | Metallothionein 1F | 2.00 | 0.751 | 0.259 | 0 |
| LINC01541 | | 1.96 | 0.979 | 0.528 | 0 |
| C2CD4B | C2 calcium-dependent domain-containing protein 4B | 1.95 | 0.607 | 0.097 | 0 |
| SCGB1D4 | Secretoglobin family 1D member 4 | 1.94 | 0.473 | 0.138 | 8.32E-194 |
| MT1H | Metallothionein 1H | 1.94 | 0.497 | 0.044 | 0 |
| CXCL14 | CXC motif chemokine 14 | 1.93 | 0.456 | 0.11 | 4.74E-238 |
| DEFB1 | Defensin beta 1 | 1.91 | 0.889 | 0.451 | 0 |
| PAPSS1 | 3'-Phosphoadenosine 5'-Phosphosulfate Synthase 1 | 1.87 | 0.939 | 0.659 | 0 |
| SLC26A2 | Sulphate transporter | 1.86 | 0.829 | 0.481 | 3.14E-237 |
| RIMKLB | Ribosomal Modification Protein RimK Like Family Member B | 1.84 | 0.925 | 0.496 | 0 |
| PAEP | Progestagen Associated Endometrial Protein | 1.82 | 0.858 | 0.642 | 2.92E-178 |
| SPP1 | Secreted phosphoprotein 1 | 1.82 | 0.741 | 0.277 | 1.35E-264 |
| MAOA | Monoamine oxidase A | 1.80 | 0.586 | 0.132 | 0 |
| MGST1 | Microsomal glutathione S-transferase 1 | 1.79 | 0.969 | 0.615 | 0 |
| SCGB2A1 | Secretoglobin family 2A member 1 | 1.78 | 0.984 | 0.769 | 0 |
| TMEM101 | Transmembrane protein 101 | 1.77 | 0.92 | 0.539 | 0 |
| SGK1 | Serum/glucocorticoid regulated kinase 1 | 1.74 | 0.947 | 0.638 | 3.88E-294 |
| ENPP3 | Ectonucleotide Pyrophosphatase/Phosphodiesterase 3 | 1.73 | 0.538 | 0.149 | 6.14E-263 |
| GABRP | Gamma-Aminobutyric Acid Type A Receptor Subunit Pi | 1.72 | 0.946 | 0.508 | 0 |
| MT1E | Metallothionein 1E | 1.72 | 0.869 | 0.555 | 4.33E-206 |
| RPS6KA5 | Ribosomal protein S6 kinase A5 | 1.71 | 0.819 | 0.347 | 0 |
| C2CD4A | C2 calcium dependent domain containing 4A | 1.71 | 0.527 | 0.048 | 0 |
| RXFP1 | Relaxin receptor 1 | 1.71 | 0.822 | 0.373 | 0 |
| FAM110C | Protein FAM110C | 1.71 | 0.541 | 0.094 | 0 |
| DEPP1 | DEPP autophagy regulator 1 | 1.69 | 0.555 | 0.191 | 2.70E-205 |
| STC1 | Stanniocalcin 1 | 1.67 | 0.585 | 0.206 | 1.10E-194 |
| SLC34A2 | Sodium-dependent phosphate transport protein 2B | 1.67 | 0.824 | 0.365 | 2.10E-286 |

**Table S8.** Top 30 genes in cluster 2.

| Gene | Protein name | Fold change | pct.1 | pct.2 | p_val_adj |
|---|---|---|---|---|---|
| TRH | Thyrotropin Releasing Hormone | 2.06 | 0.732 | 0.099 | 0 |
| CST1 | Cystatin-SN | 1.96 | 0.591 | 0.097 | 0 |
| IHH | Indian Hedgehog Signalling Molecule | 1.81 | 0.645 | 0.088 | 0 |
| SOX17 | SRY-Box Transcription Factor 17 | 1.78 | 0.952 | 0.575 | 0 |
| SOX9 | SRY-Box Transcription Factor 9 | 1.74 | 0.723 | 0.168 | 0 |
| SERPINA5 | Serpin Family A Member 5 / Protein C Inhibitor | 1.73 | 0.754 | 0.154 | 0 |
| PRR15 | Proline Rich protein 15 | 1.71 | 0.785 | 0.15 | 0 |
| CPM | Carboxypeptidase M | 1.69 | 0.812 | 0.494 | 5.09E-209 |
| SCGB2A1 | Secretoglobin Family 2A Member 1 / Mammaglobin-B | 1.68 | 0.992 | 0.77 | 2.21E-290 |
| MSX1 | Msh Homeobox 1 | 1.67 | 0.982 | 0.606 | 0 |
| CLU | Clusterin | 1.65 | 0.897 | 0.706 | 1.43E-207 |
| ASRGL1 | Asparaginase And Isoaspartyl Peptidase 1 | 1.64 | 0.995 | 0.644 | 0 |
| KIAA1324 | Endosome-Lysosome Associated Apoptosis And Autophagy Regulator 1 | 1.61 | 0.923 | 0.406 | 0 |
| CD74 | HLA class II histocompatibility antigen gamma chain | 1.60 | 0.95 | 0.656 | 1.03E-230 |
| LAMP5 | Lysosomal Associated Membrane Protein Family Member 5 | 1.59 | 0.651 | 0.064 | 0 |
| PIGR | Polymeric Immunoglobulin Receptor | 1.58 | 0.93 | 0.58 | 8.05E-228 |
| MSX2 | Msh Homeobox 2 | 1.58 | 0.652 | 0.144 | 0 |
| ID1 | DNA-binding protein inhibitor ID-1 | 1.57 | 0.985 | 0.784 | 0 |
| HLA-DRA | Major Histocompatibility Complex Class II, DR Alpha | 1.54 | 0.813 | 0.481 | 2.17E-171 |
| SMAD9 | Mothers against decapentaplegic homolog 9 | 1.54 | 0.727 | 0.189 | 0 |
| HLA-DMA | Major Histocompatibility Complex Class II DM Alpha | 1.53 | 0.791 | 0.219 | 0 |
| STXBP6 | Syntaxin Binding Protein 6 | 1.53 | 0.778 | 0.18 | 0 |
| KRT18 | Keratin type 1 cystoskeletal 18 | 1.52 | 0.999 | 0.709 | 7.25E-260 |
| DLX5 | Homeobox protein DLX-5 | 1.51 | 0.886 | 0.478 | 2.24E-244 |
| ELF3 | ETS-related transcription factor Elf-3 | 1.51 | 0.982 | 0.683 | 4.75E-207 |
| ACSL5 | Acetyl-CoA Synthetase Long Chain Family Member 5 | 1.51 | 0.953 | 0.588 | 8.77E-237 |
| PKP4 | Plakophilin 4 | 1.50 | 0.739 | 0.21 | 0 |
| STX18 | Syntaxin 18 | 1.50 | 0.941 | 0.592 | 1.18E-275 |
| MECOM | Histone-lysine N-methyltransferase MECOM | 1.50 | 0.867 | 0.485 | 3.82E-233 |
| TPD52L1 | Tumor protein D53 | 1.49 | 0.927 | 0.477 | 1.92E-257 |

**Table S9.** Top 30 genes in cluster 3.

| Gene | Protein name | Fold change | pct.1 | pct.2 | p_val_adj |
|---|---|---|---|---|---|
| C20orf85 | Ciliary microtubule inner protein 1 | 3.38 | 0.918 | 0.036 | 0 |
| RSPH1 | Radial Spoke Head Component 1 | 3.02 | 0.853 | 0.057 | 0 |
| C9orf24 | Sperm Microtubule Inner Protein 6 | 3.01 | 0.848 | 0.038 | 0 |
| C11orf88 | Cilia- and flagella-associated protein HOATZ | 2.86 | 0.821 | 0.03 | 0 |
| C1orf194 | Cilia And Flagella Associated Protein 276 | 2.84 | 0.803 | 0.038 | 0 |
| SNTN | Sentan, Cilia Apical Structure Protein | 2.81 | 0.81 | 0.025 | 0 |
| AGR3 | Anterior Gradient 3 | 2.74 | 0.816 | 0.121 | 0 |
| TPPP3 | Tubulin polymerization-promoting protein family member 3 | 2.74 | 0.936 | 0.254 | 0 |
| PIFO | Ciliary microtubule associated protein 3 | 2.68 | 0.773 | 0.045 | 0 |
| FAM183A | Cilia And Flagella Associated Protein 144 | 2.61 | 0.759 | 0.028 | 0 |
| CAPS | Calcyphosine | 2.59 | 1 | 0.622 | 0 |
| CAPSL | Calcyphosine like protein | 2.55 | 0.742 | 0.025 | 0 |
| C5orf49 | Cilia and flagella associated protein 90 | 2.52 | 0.748 | 0.042 | 0 |
| MORN2 | MORN repeat-containing protein 2 | 2.48 | 0.81 | 0.219 | 0 |
| C9orf116 | Piecer of microtubule wall 1 | 2.35 | 0.732 | 0.105 | 0 |
| LRRIQ1 | Leucine-rich repeats and IQ domain-containing protein 1 | 2.33 | 0.688 | 0.035 | 0 |
| AL357093.2 | | 2.29 | 0.662 | 0.03 | 0 |
| MS4A8 | Membrane spanning 4-domains A8 | 2.28 | 0.644 | 0.033 | 0 |
| ODF3B | Ciliary Microtubule Associated Protein 1B | 2.25 | 0.752 | 0.206 | 0 |
| MORN5 | MORN repeat-containing protein 5 | 2.24 | 0.666 | 0.024 | 0 |
| DNAAF1 | Dynein Axonemal Assembly Factor 1 | 2.24 | 0.657 | 0.028 | 0 |
| DYDC2 | DPY30 Domain Containing 2 | 2.23 | 0.647 | 0.021 | 0 |
| ZMYND10 | Zinc Finger MYND-Type Containing 10 | 2.23 | 0.641 | 0.027 | 0 |
| TMEM190 | Transmembrane protein 190 | 2.21 | 0.613 | 0.021 | 0 |
| AC007906.2 | | 2.16 | 0.664 | 0.062 | 0 |
| CFAP53 | Cilia and flagella associated protein 53 | 2.14 | 0.639 | 0.024 | 0 |
| FAM92B | CBY1 Interacting BAR Domain Containing 2 | 2.14 | 0.634 | 0.017 | 0 |
| ROPN1L | Rhophilin Associated Tail Protein 1 Like | 2.13 | 0.62 | 0.019 | 0 |
| SPA17 | Sperm Autoantigenic Protein 17 | 2.10 | 0.65 | 0.113 | 0 |
| CFAP126 | Cilia And Flagella Associated Protein 126 | 2.10 | 0.613 | 0.019 | 0 |

**Table S10.** Top 30 genes in cluster 4.

| Gene | Protein name | fold change | pct.1 | pct.2 | p_val_adj |
|---|---|---|---|---|---|
| RGS5 | Regulator of G protein signalling 5 | 2.23 | 0.591 | 0.102 | 0 |
| IGFBP5 | Insulin Like Growth Factor Binding Protein 5 | 2.13 | 0.924 | 0.542 | 2.35E-258 |
| GUCY1A2 | Guanylate Cyclase 1 Soluble Subunit Alpha 2 | 2.03 | 0.751 | 0.116 | 0 |
| LHFPL6 | LHFPL Tetraspan Subfamily Member 6 | 1.98 | 0.774 | 0.197 | 0 |
| ACTA2 | Actin Alpha 2 | 1.94 | 0.917 | 0.515 | 1.54E-222 |
| NOTCH3 | Notch receptor 3 | 1.90 | 0.795 | 0.183 | 0 |
| EBF1 | Transcription factor COE1 | 1.88 | 0.768 | 0.168 | 0 |
| COL4A1 | Collagen Type IV Alpha 1 Chain | 1.86 | 0.964 | 0.577 | 3.27E-276 |
| ZEB2 | Zinc Finger E-Box Binding Homeobox 2 | 1.83 | 0.797 | 0.234 | 0 |
| CAV1 | Caveolin 1 | 1.83 | 0.933 | 0.452 | 2.64E-271 |
| MGP | Matrix Gla Protein | 1.83 | 0.983 | 0.571 | 9.23E-230 |
| C11orf96 | Uncharacterized Protein C11orf96 | 1.82 | 0.992 | 0.626 | 8.70E-241 |
| COL14A1 | Collagen Type XIV Alpha 1 Chain | 1.79 | 0.691 | 0.202 | 7.11E-257 |
| SYNPO2 | Synaptopodin 2 | 1.78 | 0.785 | 0.249 | 3.25E-269 |
| FILIP1L | Filamin A Interacting Protein 1 Like | 1.78 | 0.704 | 0.205 | 3.08E-252 |
| SPARC | Secreted Protein Acidic And Cysteine Rich | 1.78 | 0.986 | 0.596 | 1.91E-233 |
| CPE | Carboxypeptidase E | 1.78 | 0.656 | 0.136 | 0 |
| ADAMTS4 | ADAM Metallopeptidase With Thrombospondin Type 1 Motif 4 | 1.77 | 0.627 | 0.134 | 0 |
| PI15 | Peptidase Inhibitor 15 | 1.75 | 0.498 | 0.07 | 0 |
| TAGLN | Transgelin | 1.75 | 0.956 | 0.559 | 7.01E-185 |
| CCL2 | C-C Motif Chemokine Ligand 2 | 1.74 | 0.491 | 0.151 | 3.21E-143 |
| GJA4 | Gap Junction Protein Alpha 4 | 1.73 | 0.634 | 0.158 | 8.14E-271 |
| THBS1 | Thrombospondin 1 | 1.72 | 0.841 | 0.449 | 6.75E-180 |
| ABCC9 | ATP Binding Cassette Subfamily C Member 9 | 1.72 | 0.616 | 0.088 | 0 |
| MYL9 | Myosin Light Chain 9 | 1.72 | 0.985 | 0.588 | 8.85E-224 |
| CALD1 | Caldesmon 1 | 1.69 | 1 | 0.798 | 0 |
| COL4A2 | Collagen Type IV Alpha 2 Chain | 1.69 | 0.97 | 0.628 | 3.71E-236 |
| CRISPLD2 | Cysteine-rich secretory protein LCCL domain-containing 2 | 1.68 | 0.859 | 0.39 | 3.86E-217 |
| PPP1R14A | Protein Phosphatase 1 Regulatory Inhibitor Subunit 14A | 1.66 | 0.739 | 0.23 | 1.11E-224 |
| MYLK | Myosin Light Chain Kinase | 1.65 | 0.731 | 0.249 | 1.53E-201 |

**Table S11.** Top 30 genes in cluster 5.

| Gene | Protein name | fold change | pct.1 | pct.2 | p_val_adj |
|---|---|---|---|---|---|
| MMP7 | Matrix metallopeptidase 7 | 2.22 | 0.982 | 0.612 | 0 |
| AREG | Amphiregulin | 1.90 | 0.68 | 0.165 | 2.16E-294 |
| HMGA1 | High Mobility Group AT-Hook | 1.88 | 0.972 | 0.54 | 0 |
| KRT7 | Keratin 7 | 1.87 | 0.867 | 0.341 | 5.31E-275 |
| TM4SF1 | Transmembrane 4 L6 Family Member 1 | 1.87 | 0.954 | 0.526 | 3.32E-257 |
| TACSTD2 | Tumor Associated Calcium Signal Transducer 2 | 1.86 | 0.988 | 0.513 | 0 |
| CXCL8 | CXC motif chemokine 8 | 1.83 | 0.795 | 0.371 | 4.19E-175 |
| CXCL2 | CXC motif chemokine 2 | 1.81 | 0.852 | 0.445 | 2.94E-184 |
| TFPI2 | Tissue Factor Pathway Inhibitor 2 | 1.79 | 0.967 | 0.563 | 1.67E-247 |
| PLAUR | Urokinase plasminogen activator surface receptor | 1.77 | 0.943 | 0.54 | 2.39E-240 |
| BIRC3 | Baculoviral IAP Repeat Containing 3 | 1.73 | 0.833 | 0.368 | 1.07E-219 |
| LAMC2 | Laminin Subunit Gamma 2 | 1.72 | 0.674 | 0.164 | 7.36E-287 |
| S100A10 | S100 Calcium Binding Protein A10 | 1.72 | 0.998 | 0.691 | 2.81E-264 |
| CLCF1 | Cardiotrophin Like Cytokine Factor 1 | 1.71 | 0.723 | 0.191 | 0 |
| PHLDA2 | Pleckstrin Homology Like Domain Family A Member 2 | 1.70 | 0.962 | 0.525 | 3.25E-259 |
| G0S2 | G0/G1 Switch 2 | 1.70 | 0.664 | 0.274 | 1.78E-146 |
| KRT17 | Keratin 17 | 1.69 | 0.386 | 0.074 | 1.32E-188 |
| CXCL3 | C-X-C Motif Chemokine Ligand 3 | 1.69 | 0.587 | 0.149 | 2.24E-216 |
| LAMB3 | Laminin Subunit Beta 3 | 1.69 | 0.6 | 0.135 | 6.70E-253 |
| TNFAIP3 | TNF Alpha Induced Protein 3 | 1.67 | 0.913 | 0.53 | 3.30E-195 |
| LIF | Leukemia inhibitory factor | 1.67 | 0.661 | 0.156 | 1.14E-271 |
| PRSS22 | Brain-specific serine protease 4 | 1.64 | 0.696 | 0.147 | 0 |
| KRT19 | Keratin 19 | 1.63 | 0.988 | 0.735 | 5.88E-238 |
| DUSP5 | Dual Specificity Phosphatase 5 | 1.63 | 0.68 | 0.229 | 1.70E-205 |
| NEDD9 | Neural precursor cell expressed developmentally down-regulated 9 | 1.61 | 0.944 | 0.52 | 7.32E-239 |
| SFN | Stratifin | 1.61 | 0.686 | 0.21 | 1.04E-213 |
| PMAIP1 | Phorbol-12-Myristate-13-Acetate-Induced Protein 1 | 1.61 | 0.927 | 0.482 | 2.57E-230 |
| EPHA2 | Ephrin type-A receptor 2 | 1.60 | 0.712 | 0.175 | 0 |
| KLF5 | Krueppel-like factor 5 | 1.59 | 0.926 | 0.468 | 1.14E-237 |
| EZR | Ezrin | 1.58 | 0.987 | 0.698 | 1.34E-220 |

**Table S12**. Top 30 genes in cluster 6.

| Gene | Protein name | fold change | pct.1 | pct.2 | p_val_adj |
|---|---|---|---|---|---|
| MUSTN1 | Musculoskeletal embryonic nuclear protein 1 | 3.16 | 0.939 | 0.066 | 0 |
| ADIRF | Adipogenesis Regulatory Factor | 2.89 | 0.989 | 0.476 | 0 |
| MYH11 | Myosin Heavy Chain 11 | 2.84 | 0.97 | 0.142 | 0 |
| MCAM | Melanoma Cell Adhesion Molecule | 2.82 | 0.971 | 0.141 | 0 |
| ACTA2 | Actin, Aortic Smooth Muscle | 2.59 | 0.998 | 0.517 | 0 |
| S100A4 | S100 Calcium Binding Protein A4 | 2.45 | 0.983 | 0.508 | 0 |
| TAGLN | Transgelin | 2.40 | 1 | 0.563 | 0 |
| MYL9 | Myosin Light Chain 9 | 2.38 | 1 | 0.595 | 0 |
| PLN | Phospholamban | 2.36 | 0.852 | 0.069 | 0 |
| NOTCH3 | Notch Receptor 3 | 2.34 | 0.961 | 0.183 | 0 |
| MEF2C | Myocyte Enhancer Factor 2C | 2.34 | 0.902 | 0.118 | 0 |
| RGS5 | Regulator Of G Protein Signalling 5 | 2.27 | 0.77 | 0.099 | 0 |
| HOPX | HOP Homeobox | 2.15 | 0.785 | 0.088 | 0 |
| NDUFA4L2 | NADH dehydrogenase [ubiquinone] 1 alpha subcomplex subunit 4-like 2 | 2.15 | 0.696 | 0.055 | 0 |
| CAV1 | Caveolin 1 | 2.15 | 0.991 | 0.457 | 0 |
| SOD3 | Superoxide Dismutase 3 | 2.14 | 0.985 | 0.54 | 0 |
| PTP4A3 | Protein Tyrosine Phosphatase 4A3 | 2.12 | 0.835 | 0.095 | 0 |
| MYLK | Myosin Light Chain Kinase | 2.11 | 0.914 | 0.245 | 0 |
| SLIT3 | Slit Guidance Ligand 3 | 2.10 | 0.877 | 0.138 | 0 |
| TPM2 | Tropomyosin 2 | 2.10 | 1 | 0.623 | 0 |
| MT1M | Metallothionein 1M | 2.09 | 0.738 | 0.179 | 1.29E-277 |
| PPP1R14A | Protein Phosphatase 1 Regulatory Inhibitor Subunit 14A | 2.08 | 0.915 | 0.227 | 0 |
| TINAGL1 | Tubulointerstitial Nephritis Antigen Like 1 | 2.08 | 0.911 | 0.224 | 0 |
| FRZB | Frizzled Related Protein | 2.08 | 0.808 | 0.078 | 0 |
| NTRK2 | Neurotrophic Receptor Tyrosine Kinase 2 | 2.07 | 0.769 | 0.023 | 0 |
| TIMP3 | TIMP Metallopeptidase Inhibitor 3 | 2.06 | 0.991 | 0.537 | 4.61E-299 |
| CAVIN3 | Caveolae Associated Protein 3 | 2.06 | 0.893 | 0.213 | 0 |
| CPE | Carboxypeptidase E | 2.00 | 0.832 | 0.134 | 0 |
| PDGFA | Platelet Derived Growth Factor Subunit A | 2.00 | 0.828 | 0.192 | 0 |
| MAP3K7CL | MAP3K7 C-Terminal Like | 2.00 | 0.722 | 0.032 | 0 |

**Table S13.** Top 30 genes in cluster 7.

| Gene | Protein name | fold change | pct.1 | pct.2 | p_val_adj |
|---|---|---|---|---|---|
| CPM | Carboxypeptidase M | 1.74 | 0.951 | 0.502 | 2.67E-170 |
| SOX17 | SRY-Box Transcription Factor 17 | 1.50 | 0.975 | 0.594 | 3.71E-111 |
| DUSP2 | Dual specificity protein phosphatase 2 | 1.49 | 0.957 | 0.574 | 4.35E-102 |
| MSX2 | Homeobox protein MSX-2 | 1.43 | 0.472 | 0.184 | 6.51E-80 |
| GALNT4 | Polypeptide N-Acetylgalactosaminyltransferase 4 | 1.42 | 0.884 | 0.451 | 3.00E-110 |
| CXADR | Coxsackievirus and adenovirus receptor | 1.41 | 0.925 | 0.497 | 1.44E-112 |
| MECOM | Histone-lysine N-methyltransferase MECOM | 1.40 | 0.93 | 0.502 | 9.05E-104 |
| MMP7 | Matrix Metallopeptidase 7 | 1.40 | 0.97 | 0.621 | 4.12E-72 |
| TRH | Thyrotropin Releasing Hormone | 1.39 | 0.469 | 0.151 | 1.58E-83 |
| ASRGL1 | Asparaginase and Isoaspartyl peptidase 1 | 1.38 | 0.99 | 0.664 | 3.98E-82 |
| STX18 | Syntaxin-18 | 1.37 | 0.97 | 0.609 | 4.85E-97 |
| ID1 | DNA-Binding Protein inhibitor ID-1 | 1.37 | 0.998 | 0.794 | 4.39E-87 |
| IHH | Indian Hedgehog signalling molecule | 1.37 | 0.418 | 0.134 | 6.73E-80 |
| SPRY1 | Sprouty RTK signalling anagonist 1 | 1.36 | 0.813 | 0.444 | 3.06E-75 |
| PLCB1 | Phospholipase C-beta-1 | 1.35 | 0.884 | 0.482 | 7.27E-89 |
| CCND1 | Cyclin D1 | 1.34 | 0.948 | 0.58 | 8.46E-79 |
| GREM2 | Gremlin-2 | 1.34 | 0.411 | 0.115 | 2.54E-101 |
| SCGB2A1 | Secretoglobin Family 2A Member 1 / Mammaglobin-B | 1.34 | 0.995 | 0.782 | 2.47E-67 |
| TPD52L1 | Tumor protein D53 | 1.34 | 0.957 | 0.5 | 1.83E-96 |
| C2orf88 | Small membrane A-kinase anchor protein | 1.32 | 0.931 | 0.486 | 2.49E-86 |
| OCIAD2 | OCIA domain-containing protein 2 | 1.32 | 0.982 | 0.629 | 4.97E-76 |
| KIAA1324 | Endosome-Lysosome Associated Apoptosis and Autophagy Regulator 1 | 1.32 | 0.859 | 0.438 | 4.25E-78 |
| KRT8 | Keratin type II cytoskeletal 8 | 1.32 | 1 | 0.713 | 4.26E-65 |
| ATP1B1 | Sodium/potassium-transporting ATPase subunit beta-1 | 1.31 | 0.974 | 0.651 | 4.82E-61 |
| HPGD | 15-Hydroxyprostaglandin Dehydrogenase | 1.31 | 0.41 | 0.15 | 1.94E-64 |
| GCLC | Glutamate-Cysteine Ligase Catalytic Subunit | 1.31 | 0.441 | 0.211 | 5.07E-48 |
| PKP4 | Plakophilin 4 | 1.30 | 0.497 | 0.255 | 3.63E-49 |
| CMTM6 | CKLF-like MARVEL transmembrane domain-containing protein 6 | 1.30 | 0.982 | 0.804 | 8.51E-77 |

| | | | | | |
|---|---|---|---|---|---|
| KRT18 | Keratin-18 | 1.29 | 0.998 | 0.725 | 2.60E-52 |
| PRR15 | Proline-rich protein 15 | 1.29 | 0.444 | 0.208 | 2.15E-46 |

**Table S14.** Top 30 genes in cluster 8.

| Gene | Protein name | fold change | pct.1 | pct.2 | p_val_adj |
|---|---|---|---|---|---|
| LUM | Lumican | 2.50 | 1 | 0.527 | 1.07E-298 |
| INHBA | Inhibin Subunit Beta A | 2.49 | 0.673 | 0.095 | 0 |
| MMP10 | Matrix Metallopeptidase 10 | 2.37 | 0.443 | 0.058 | 2.43E-255 |
| COL6A3 | Collagen type VI alpha 3 chain | 2.31 | 0.998 | 0.464 | 0 |
| SPON2 | Spondin 2 | 2.13 | 0.796 | 0.239 | 2.13E-255 |
| COL1A1 | Collagen type I alpha 1 chain | 2.07 | 0.997 | 0.617 | 7.17E-242 |
| WNT5A | Wnt Family Member 5A | 2.05 | 0.742 | 0.195 | 2.96E-271 |
| SERPINE1 | Endothelial Plasminogen Activator Inhibitor | 2.04 | 0.734 | 0.188 | 1.47E-236 |
| MMP3 | Matrix metallopeptidase 3 | 2.03 | 0.351 | 0.03 | 2.11E-270 |
| COL3A1 | Collagen type III alpha 1 chain | 2.01 | 0.998 | 0.612 | 5.53E-217 |
| MMP2 | Matrix metallopeptidase 2 | 2.00 | 0.981 | 0.5 | 3.16E-253 |
| DCN | Decorin | 1.98 | 0.988 | 0.524 | 6.24E-205 |
| LOXL2 | Lysyl Oxidase Like 2 | 1.97 | 0.701 | 0.098 | 0 |
| APOE | Apolipoprotein E | 1.96 | 0.971 | 0.492 | 1.93E-199 |
| COL5A2 | Collagen Type V Alpha 2 Chain | 1.96 | 0.82 | 0.252 | 1.16E-249 |
| IGFBP6 | Insulin Like Growth Factor Binding Protein 6 | 1.91 | 0.692 | 0.22 | 6.45E-181 |
| CTHRC1 | Collagen Triple Helix Repeat Containing 1 | 1.85 | 0.569 | 0.106 | 3.72E-253 |
| ISLR | Immunoglobulin Superfamily Containing Leucine Rich Repeat | 1.84 | 0.777 | 0.241 | 2.06E-226 |
| COL6A2 | Collagen Type VI Alpha 2 Chain | 1.82 | 0.995 | 0.58 | 1.58E-204 |
| COL7A1 | Collagen Type VII Alpha 1 Chain | 1.82 | 0.625 | 0.092 | 0 |
| SPARC | Secreted Protein Acidic And Cysteine Rich | 1.81 | 0.997 | 0.607 | 1.16E-179 |
| TNFRSF11B | Tumor necrosis factor receptor superfamily member 11B | 1.80 | 0.424 | 0.043 | 9.66E-294 |
| IL11 | Interleukin 11 | 1.80 | 0.455 | 0.062 | 5.46E-249 |
| TMEM158 | Transmembrane Protein 158 | 1.78 | 0.628 | 0.143 | 2.17E-223 |
| PCOLCE | Procollagen C-Endopeptidase Enhancer | 1.77 | 0.971 | 0.511 | 4.22E-180 |
| PLAT | Tissue-Type Plasminogen Activator | 1.77 | 0.694 | 0.201 | 1.68E-195 |
| COL6A1 | Collagen Type VI Alpha 1 Chain | 1.77 | 0.984 | 0.553 | 5.26E-191 |
| LGALS1 | Galectin 1 | 1.77 | 0.998 | 0.693 | 4.92E-213 |
| STC1 | Stanniocalcin 1 | 1.76 | 0.538 | 0.233 | 2.89E-79 |
| CTSK | Cathepsin K | 1.76 | 0.713 | 0.209 | 1.15E-198 |

**Table S15.** Top 30 genes in cluster 9.

| Gene | Protein name | fold change | pct.1 | pct.2 | p_val_adj |
|---|---|---|---|---|---|
| CCL20 | C-C Motif Chemokine Ligand 20 | 3.35 | 0.808 | 0.108 | 0 |
| G0S2 | G0/G1 Switch 2 | 3.09 | 0.963 | 0.272 | 0 |
| SPP1 | Secreted Phosphoprotein 1 | 2.93 | 0.963 | 0.299 | 0 |
| PAEP | Progestagen Associated Endometrial Protein | 2.73 | 1 | 0.651 | 1.08E-272 |
| IL17C | Interleukin 17C | 2.58 | 0.654 | 0.022 | 0 |
| PTGS2 | Prostaglandin-Endoperoxide Synthase 2 | 2.52 | 0.722 | 0.134 | 0 |
| CXCL8 | C-X-C Motif Chemokine Ligand 8 | 2.38 | 0.975 | 0.376 | 1.23E-236 |
| CXCL2 | C-X-C Motif Chemokine Ligand 2 | 2.36 | 0.99 | 0.452 | 5.07E-233 |
| TFPI2 | Tissue Factor Pathway Inhibitor 2 | 2.34 | 0.994 | 0.576 | 7.84E-241 |
| SERPINA1 | Alpha-1-antitrypsin | 2.33 | 0.992 | 0.5 | 7.98E-245 |
| LYPD3 | LY6/PLAUR Domain Containing 3 | 2.26 | 0.611 | 0.053 | 0 |
| LAMB3 | Laminin Subunit Beta 3 | 2.26 | 0.722 | 0.145 | 6.08E-288 |
| VNN1 | Vanin 1 | 2.19 | 0.603 | 0.054 | 0 |
| LCN2 | Lipocalin 2 | 2.03 | 0.947 | 0.388 | 2.74E-207 |
| CYP24A1 | Cytochrome P450 Family 24 Subfamily A Member 1 | 2.00 | 0.605 | 0.113 | 1.07E-236 |
| AQP3 | Aquaporin-3 | 1.99 | 0.941 | 0.406 | 2.67E-207 |
| SMOX | Spermine Oxidase | 1.99 | 0.62 | 0.155 | 2.41E-201 |
| SLPI | Secretory Leukocyte Peptidase Inhibitor | 1.98 | 1 | 0.712 | 4.52E-221 |
| CLDN10 | Claudin 10 | 1.98 | 0.971 | 0.45 | 5.58E-213 |
| TNFAIP2 | TNF Alpha Induced Protein 2 | 1.98 | 0.982 | 0.492 | 6.10E-192 |
| GPX3 | Glutathione Peroxidase 3 | 1.96 | 0.996 | 0.481 | 8.04E-192 |
| C15orf48 | Normal mucosa of esophagus-specific gene 1 protein | 1.94 | 0.581 | 0.135 | 2.17E-185 |
| PDZK1IP1 | PDZK1 Interacting Protein 1 | 1.90 | 0.654 | 0.165 | 1.62E-206 |
| DUSP4 | Dual Specificity Phosphatase 4 | 1.89 | 0.638 | 0.192 | 2.26E-167 |
| ITGB6 | Integrin Subunit Beta 6 | 1.87 | 0.573 | 0.069 | 0 |
| SOD2 | Superoxide Dismutase 2 | 1.87 | 1 | 0.865 | 9.94E-254 |
| PLAUR | Urokinase plasminogen activator surface receptor | 1.86 | 0.994 | 0.552 | 1.86E-181 |
| MMP7 | Matrix metallopeptidase 7 | 1.85 | 0.984 | 0.625 | 2.42E-145 |
| LUCAT1 | Lung Cancer Associated Transcript 1 | 1.80 | 0.509 | 0.046 | 0 |
| LAMC2 | Laminin Subunit Gamma 2 | 1.80 | 0.616 | 0.186 | 5.90E-149 |

**Table S16.** Top 30 genes in cluster 10.

| Gene | Protein name | fold change | pct.1 | pct.2 | p_val_adj |
|---|---|---|---|---|---|
| DES | Desmin | 3.50 | 0.93 | 0.088 | 0 |
| ACTG2 | Actin Gamma 2, Smooth Muscle | 3.38 | 0.913 | 0.114 | 0 |
| CNN1 | Calponin 1 | 3.17 | 0.969 | 0.179 | 0 |
| MYH11 | Myosin Heavy Chain 11 | 2.89 | 0.941 | 0.176 | 2.46E-273 |
| MYLK | Myosin light chain kinase | 2.64 | 0.951 | 0.271 | 6.67E-216 |
| ACTA2 | Actin Alpha 2, Smooth Muscle | 2.60 | 1 | 0.536 | 1.73E-158 |
| TAGLN | Transgelin | 2.51 | 1 | 0.581 | 5.50E-158 |
| PDLIM3 | PDZ And LIM Domain 3 | 2.47 | 0.829 | 0.118 | 1.21E-291 |
| CARMN | | 2.42 | 0.808 | 0.155 | 2.10E-221 |
| KCNMA1 | Potassium Calcium-Activated Channel Subfamily M Alpha 1 | 2.35 | 0.864 | 0.13 | 0 |
| RAMP1 | Receptor Activity Modifying Protein 1 | 2.21 | 0.983 | 0.462 | 2.65E-163 |
| FHL1 | Four And A Half LIM Domains 1 | 2.17 | 0.857 | 0.199 | 6.33E-197 |
| SLMAP | Sarcolemma Associated Protein | 2.17 | 0.815 | 0.269 | 2.55E-150 |
| TPM2 | Tropomyosin 2 | 2.13 | 1 | 0.639 | 2.54E-142 |
| SFRP4 | Secreted Frizzled Related Protein 4 | 2.12 | 0.846 | 0.201 | 5.50E-160 |
| CSRP1 | Cysteine And Glycine Rich Protein 1 | 2.10 | 0.993 | 0.638 | 1.36E-153 |
| MYL9 | Myosin Light Chain 9 | 2.10 | 0.997 | 0.611 | 1.92E-125 |
| TPM1 | Tropomyosin 1 | 2.10 | 1 | 0.763 | 1.63E-153 |
| PPP1R12B | Protein Phosphatase 1 Regulatory Subunit 12B | 2.09 | 0.846 | 0.204 | 5.74E-187 |
| RSPO3 | R-Spondin 3 | 2.08 | 0.703 | 0.045 | 0 |
| FILIP1L | Filamin A Interacting Protein 1 Like | 2.06 | 0.822 | 0.231 | 7.39E-144 |
| SPARCL1 | SPARC Like 1 | 2.06 | 1 | 0.644 | 2.29E-134 |
| SVIL | Supervillin | 2.05 | 0.881 | 0.233 | 1.06E-194 |
| SYNPO2 | Synaptopodin 2 | 2.04 | 0.881 | 0.278 | 2.18E-145 |
| COL12A1 | Collagen Type XII Alpha 1 Chain | 2.04 | 0.797 | 0.216 | 6.69E-158 |
| SMTN | Smoothelin | 1.99 | 0.829 | 0.209 | 1.96E-181 |
| SMOC2 | SPARC-related modular calcium-binding protein 2 | 1.98 | 0.78 | 0.183 | 4.64E-177 |
| LMOD1 | Leiomodin 1 | 1.98 | 0.871 | 0.193 | 6.81E-198 |
| CXCL12 | C-X-C Motif Chemokine Ligand 12 | 1.93 | 0.759 | 0.142 | 6.33E-193 |
| NEXN | Nexilin F-Actin Binding Protein | 1.88 | 0.741 | 0.118 | 3.24E-222 |

**Table S17.** Top 30 genes in cluster 11.

| Gene | Protein name | fold change | pct.1 | pct.2 | p_val_adj |
|---|---|---|---|---|---|
| SAA1 | Serum Amyloid A1 | 2.96 | 0.849 | 0.08 | 0 |
| MUC5B | Mucin 5B | 2.56 | 0.773 | 0.062 | 0 |
| LCN2 | Lipocalin 2 | 2.47 | 0.945 | 0.403 | 2.06E-125 |
| SAA2 | Serum Amyloid A2 | 2.44 | 0.765 | 0.06 | 0 |
| REG1A | Regenerating Family Member 1 Alpha | 2.29 | 0.584 | 0.011 | 0 |
| CXCL5 | C-X-C Motif Chemokine Ligand 5 | 2.16 | 0.576 | 0.028 | 0 |
| CFB | Complement Factor B | 2.12 | 0.765 | 0.234 | 6.65E-110 |
| C3 | Complement C3 | 2.01 | 0.916 | 0.428 | 4.64E-102 |
| TNFRSF6B | TNF Receptor Superfamily Member 6b | 1.92 | 0.647 | 0.08 | 5.37E-202 |
| HLA-DRB5 | Major Histocompatibility Complex, Class II, DR Beta 5 | 1.89 | 0.744 | 0.21 | 2.75E-102 |
| HLA-DPA1 | Major Histocompatibility Complex, Class II, DP Alpha 1 | 1.86 | 0.693 | 0.141 | 1.87E-126 |
| LTF | Lactotransferrin | 1.84 | 0.496 | 0.025 | 0 |
| HLA-DRA | Major Histocompatibility Complex, Class II, DR Alpha | 1.82 | 0.95 | 0.508 | 1.40E-78 |
| CEACAM5 | Carcinoembryonic Antigen-Related Cell Adhesion Molecule 5 | 1.81 | 0.378 | 0.004 | 0 |
| HLA-DQA1 | Major Histocompatibility Complex, Class II, DQ Alpha 1 | 1.80 | 0.571 | 0.054 | 5.31E-218 |
| MSLN | Mesothelin | 1.80 | 0.479 | 0.039 | 6.75E-215 |
| CCL20 | C-C Motif Chemokine Ligand 20 | 1.74 | 0.513 | 0.134 | 1.01E-58 |
| CD74 | CD74 Molecule | 1.73 | 0.992 | 0.682 | 2.45E-77 |
| IDO1 | Indoleamine 2,3-Dioxygenase 1 | 1.73 | 0.655 | 0.219 | 4.02E-64 |
| HLA-DRB1 | Major Histocompatibility Complex, Class II, DR Beta 1 | 1.72 | 0.882 | 0.424 | 1.82E-69 |
| SLPI | Secretory Leukocyte Peptidase Inhibitor | 1.68 | 0.992 | 0.72 | 6.77E-66 |
| FGFBP1 | Fibroblast Growth Factor Binding Protein 1 | 1.68 | 0.387 | 0.031 | 1.20E-169 |
| HLA-DPB1 | Major Histocompatibility Complex, Class II, DP Beta 1 | 1.64 | 0.592 | 0.184 | 2.54E-67 |
| MUC1 | Mucin-1 | 1.60 | 0.916 | 0.64 | 1.84E-49 |
| S100A4 | S100 Calcium Binding Protein A4 | 1.60 | 0.954 | 0.53 | 5.53E-52 |
| PIGR | Polymeric Immunoglobulin Receptor | 1.60 | 0.929 | 0.612 | 3.31E-48 |
| ADIRF | Adipogenesis Regulatory Factor | 1.60 | 0.933 | 0.501 | 1.36E-49 |
| AGR2 | Anterior Gradient 2, Protein Disulphide Isomerase Family Member | 1.60 | 0.849 | 0.495 | 1.02E-45 |
| CSF3 | Colony Stimulating Factor 3 | 1.57 | 0.5 | 0.115 | 1.23E-69 |
| FXYD3 | FXYD Domain Containing Ion Transport Regulator 3 | 1.57 | 0.874 | 0.43 | 1.16E-59 |

**Table S18.** Top 30 genes in cluster 12.

| Gene | Protein name | fold change | pct.1 | pct.2 | p_val_adj |
|---|---|---|---|---|---|
| CCDC80 | Coiled-Coil Domain Containing 80 | 3.02 | 0.961 | 0.256 | 3.66E-172 |
| OGN | Osteoglycin | 2.97 | 0.877 | 0.13 | 1.92E-245 |
| DCN | Decorin | 2.50 | 1 | 0.542 | 6.56E-121 |
| C7 | Complement 7 | 2.30 | 0.581 | 0.016 | 0 |
| ASPN | Asporin | 2.22 | 0.768 | 0.096 | 4.15E-218 |
| MGP | Matrix Gla Protein | 2.19 | 0.99 | 0.599 | 6.27E-88 |
| LUM | Lumican | 2.18 | 1 | 0.545 | 1.13E-89 |
| CXCL12 | C-X-C Motif Chemokine Ligand 12 | 2.18 | 0.862 | 0.145 | 3.44E-194 |
| DPT | Dermatopontin | 2.16 | 0.7 | 0.055 | 3.84E-300 |
| SERPINF1 | Pigment epithelium-derived factor | 2.12 | 0.956 | 0.397 | 1.11E-102 |
| SPARCL1 | SPARC-like protein 1 | 2.12 | 1 | 0.648 | 6.85E-102 |
| APOD | Apolipoprotein D | 2.11 | 0.729 | 0.145 | 6.76E-122 |
| CTSK | Cathepsin K | 2.08 | 0.862 | 0.226 | 6.07E-129 |
| SFRP1 | Secreted Frizzled Related Protein 1 | 2.08 | 0.788 | 0.155 | 7.30E-141 |
| PLAC9 | Placenta Associated 9 | 2.03 | 0.833 | 0.201 | 2.51E-126 |
| SCN7A | Sodium Voltage-Gated Channel Alpha Subunit 7 | 2.03 | 0.512 | 0.005 | 0 |
| MEG3 | Maternally Expressed 3 | 2.01 | 0.897 | 0.232 | 6.07E-111 |
| COL14A1 | Collagen Type XIV Alpha 1 Chain | 1.98 | 0.818 | 0.232 | 1.58E-100 |
| IGFBP5 | Insulin Like Growth Factor Binding Protein 5 | 1.98 | 0.901 | 0.568 | 5.22E-55 |
| GPNMB | Transmembrane glycoprotein NMB | 1.97 | 0.823 | 0.193 | 4.62E-132 |
| BGN | Biglycan | 1.97 | 0.931 | 0.415 | 4.54E-87 |
| IGFBP6 | Insulin Like Growth Factor Binding Protein 6 | 1.96 | 0.803 | 0.236 | 5.71E-92 |
| IGF1 | Insulin Like Growth Factor 1 | 1.95 | 0.877 | 0.222 | 2.33E-99 |
| SPON2 | Spondin 2 | 1.92 | 0.887 | 0.26 | 8.79E-100 |
| FRZB | Frizzled Related Protein | 1.91 | 0.714 | 0.115 | 3.65E-146 |
| TIMP2 | TIMP Metallopeptidase Inhibitor 2 | 1.90 | 0.921 | 0.322 | 1.85E-101 |
| SPOCK1 | Testican-1 | 1.90 | 0.729 | 0.113 | 1.19E-164 |
| AEBP1 | Adipocyte enhancer-binding protein 1 | 1.88 | 0.901 | 0.293 | 1.15E-99 |
| COL6A2 | Collagen Type VI Alpha 2 Chain | 1.87 | 0.99 | 0.597 | 2.60E-81 |
| A2M | Alpha-2-Macroglobulin | 1.87 | 0.837 | 0.235 | 7.69E-100 |

**Table S19.** Top 30 genes in cluster 13.

| Gene | Protein name | fold change | pct.1 | pct.2 | p_val_adj |
|---|---|---|---|---|---|
| PCLAF | PCNA Clamp Associated Factor | 2.11 | 0.602 | 0.076 | 6.59E-151 |
| RRM2 | Ribonucleotide Reductase Regulatory Subunit M2 | 1.78 | 0.541 | 0.027 | 6.66E-296 |
| TYMS | Thymidylate Synthetase | 1.70 | 0.569 | 0.055 | 2.12E-175 |
| TK1 | Thymidine Kinase 1 | 1.70 | 0.552 | 0.057 | 9.38E-162 |
| HMGB2 | High Mobility Group Box 2 | 1.66 | 0.652 | 0.304 | 5.21E-40 |
| UBE2T | Ubiquitin-conjugating enzyme E2 T | 1.64 | 0.541 | 0.08 | 1.25E-112 |
| CDK1 | Cyclin Dependent Kinase 1 | 1.64 | 0.519 | 0.049 | 1.35E-161 |
| PCNA | Proliferating Cell Nuclear Antigen | 1.63 | 0.591 | 0.224 | 5.65E-43 |
| ZWINT | ZW10 Interacting Kinetochore Protein | 1.63 | 0.519 | 0.056 | 1.65E-146 |
| UBE2C | Ubiquitin Conjugating Enzyme E2 C | 1.61 | 0.508 | 0.022 | 1.45E-301 |
| HELLS | Lymphoid-specific helicase | 1.61 | 0.514 | 0.136 | 2.09E-55 |
| HIST1H4C | H4 Clustered Histone 3 | 1.60 | 0.867 | 0.597 | 1.57E-34 |
| AREG | Amphiregulin | 1.57 | 0.558 | 0.202 | 8.22E-31 |
| H2AFZ | H2A.Z Variant Histone 1 | 1.56 | 0.972 | 0.888 | 1.94E-52 |
| MCM7 | DNA replication licensing factor MCM7 | 1.56 | 0.547 | 0.179 | 1.49E-46 |
| CLSPN | Claspin | 1.55 | 0.47 | 0.031 | 6.98E-203 |
| NUSAP1 | Nucleolar And Spindle Associated Protein 1 | 1.55 | 0.464 | 0.028 | 4.54E-211 |
| TMEM106C | Transmembrane Protein 106C | 1.55 | 0.58 | 0.241 | 2.23E-37 |
| MAD2L1 | Mitotic Arrest Deficient 2 Like 1 | 1.55 | 0.508 | 0.073 | 1.27E-105 |
| TOP2A | DNA Topoisomerase II Alpha | 1.54 | 0.409 | 0.024 | 1.76E-186 |
| RRM1 | Ribonucleotide Reductase Catalytic Subunit M1 | 1.53 | 0.547 | 0.136 | 2.23E-64 |
| MCM3 | DNA replication licensing factor MCM3 | 1.52 | 0.547 | 0.166 | 2.94E-49 |
| CENPF | Centromere Protein F | 1.52 | 0.436 | 0.046 | 5.01E-117 |
| DHFR | Dihydrofolate Reductase | 1.51 | 0.503 | 0.083 | 7.96E-89 |
| CKS1B | Cyclin-dependent kinases regulatory subunit 1 | 1.51 | 0.558 | 0.219 | 3.10E-37 |
| DUSP2 | Dual Specificity Phosphatase 2 | 1.50 | 0.945 | 0.592 | 1.43E-30 |
| BIRC5 | Baculoviral IAP Repeat Containing 5 | 1.48 | 0.431 | 0.021 | 4.61E-236 |
| STMN1 | Stathmin 1 | 1.48 | 0.956 | 0.748 | 1.96E-34 |
| CENPU | Centromere Protein U | 1.48 | 0.464 | 0.043 | 1.96E-144 |
| LMNB1 | Lamin B1 | 1.48 | 0.492 | 0.053 | 4.48E-134 |

**Table S20.** Top 30 genes in cluster 14.

| Gene | Protein name | fold change | pct.1 | pct.2 | p_val_adj |
|---|---|---|---|---|---|
| MT1G | Metallothionein 1G | 1.92 | 0.847 | 0.443 | 1.91E-15 |
| SERPINA5 | Plasma serine protease inhibitor | 1.85 | 0.671 | 0.218 | 1.14E-26 |
| MT1F | Metallothionein 1F | 1.76 | 0.729 | 0.315 | 5.46E-19 |
| TMEM101 | Transmembrane Protein 101 | 1.72 | 0.976 | 0.581 | 1.01E-21 |
| SCGB1D2 | Secretoglobin family 1D member 2 | 1.71 | 0.871 | 0.54 | 3.40E-11 |
| PTGS1 | Prostaglandin-Endoperoxide Synthase 1 | 1.71 | 0.459 | 0.037 | 8.72E-85 |
| MT1E | Metallothionein 1E | 1.68 | 0.929 | 0.59 | 9.08E-15 |
| SLC26A2 | Solute Carrier Family 26 Member 2 | 1.65 | 0.941 | 0.519 | 8.15E-18 |
| MAP2K6 | Mitogen-Activated Protein Kinase Kinase 6 | 1.64 | 0.635 | 0.234 | 1.21E-19 |
| PSAT1 | Phosphoserine Aminotransferase 1 | 1.63 | 0.529 | 0.129 | 1.07E-27 |
| AC025580.1 |  | 1.63 | 0.494 | 0.128 | 5.74E-23 |
| ENPP3 | Ectonucleotide Pyrophosphatase/Phosphodiesterase 3 | 1.60 | 0.506 | 0.193 | 2.92E-12 |
| UCA1 | Urothelial Cancer Associated 1 | 1.59 | 0.976 | 0.626 | 4.24E-18 |
| PKHD1L1 | Fibrocystin-L | 1.59 | 0.565 | 0.138 | 4.34E-29 |
| MT1H | Metallothionein 1H | 1.58 | 0.4 | 0.096 | 1.26E-17 |
| RNASET2 | Ribonuclease T2 | 1.57 | 1 | 0.727 | 8.64E-21 |
| HMGCR | 3-Hydroxy-3-Methylglutaryl-CoA Reductase | 1.55 | 0.624 | 0.269 | 5.28E-15 |
| FAM110C | Protein FAM110C | 1.54 | 0.518 | 0.144 | 6.01E-20 |
| SLC39A14 | Metal cation symporter ZIP14 | 1.53 | 0.459 | 0.241 | 3.07E-05 |
| ASRGL1 | Asparaginase And Isoaspartyl Peptidase 1 | 1.53 | 1 | 0.682 | 3.20E-19 |
| PAX2 | Paired Box 2 | 1.51 | 0.471 | 0.199 | 3.95E-10 |
| C2orf88 | Small membrane A-kinase anchor protein | 1.50 | 0.976 | 0.511 | 1.11E-20 |
| PLLP |  | 1.50 | 0.624 | 0.214 | 6.30E-20 |
| MT1X | Metallothionein 1X | 1.50 | 0.906 | 0.579 | 1.31E-10 |
| MUC1 | Mucin-1 | 1.50 | 1 | 0.644 | 5.21E-17 |
| LINC01541 |  | 1.48 | 0.941 | 0.579 | 1.64E-14 |
| MSX1 | Msh Homeobox 1 | 1.48 | 1 | 0.646 | 1.31E-17 |
| MPZL2 | Myelin Protein Zero Like 2 | 1.48 | 0.941 | 0.49 | 4.21E-18 |
| FGF9 | Fibroblast Growth Factor 9 | 1.48 | 0.412 | 0.073 | 7.43E-30 |
| SH3YL1 | SH3 And SYLF Domain Containing 1 | 1.47 | 0.965 | 0.592 | 9.86E-17 |

**Table S21.** Top 30 genes in cluster 15.

| Gene | Protein name | fold change | pct.1 | pct.2 | p_val_adj |
|---|---|---|---|---|---|
| SAA1 | Serum Amyloid A1 | 3.01 | 0.86 | 0.095 | 2.43E-73 |
| C20orf85 | Ciliary Microtubule Inner Protein 1 | 2.83 | 1 | 0.114 | 3.01E-80 |
| C9orf24 | Sperm Microtubule Inner Protein 6 | 2.58 | 1 | 0.109 | 3.20E-81 |
| C1orf194 | Cilia And Flagella Associated Protein 276 | 2.55 | 0.96 | 0.105 | 9.09E-80 |
| MMP3 | Matrix metallopeptidase 3 | 2.50 | 0.62 | 0.046 | 4.33E-77 |
| AGR3 | Anterior Gradient 3 | 2.44 | 0.94 | 0.182 | 8.65E-47 |
| C11orf88 | HOATZ Cilia And Flagella Associated Protein | 2.39 | 0.92 | 0.1 | 7.19E-75 |
| FAM183A | Cilia And Flagella Associated Protein 144 | 2.33 | 0.88 | 0.093 | 5.76E-74 |
| SAA2 | Serum Amyloid A2 | 2.32 | 0.7 | 0.074 | 1.45E-58 |
| RSPH1 | Radial Spoke Head Component 1 | 2.30 | 0.9 | 0.128 | 1.48E-54 |
| TPPP3 | Tubulin Polymerization Promoting Protein Family Member 3 | 2.28 | 0.96 | 0.314 | 7.36E-30 |
| CAPS | Nuclear Cap Binding Protein Subunit 1 | 2.27 | 1 | 0.656 | 4.33E-24 |
| CAPSL | Calcyphosine-like | 2.25 | 0.88 | 0.088 | 1.59E-76 |
| MORN2 | MORN Repeat Containing 2 | 2.16 | 0.94 | 0.271 | 1.26E-32 |
| PIFO | Ciliary Microtubule Associated Protein 3 | 2.16 | 0.84 | 0.11 | 9.35E-55 |
| RRAD | GTP-binding protein RAD | 2.12 | 0.9 | 0.21 | 2.30E-36 |
| AL357093.2 | | 2.12 | 0.84 | 0.086 | 4.32E-72 |
| MS4A8 | Membrane Spanning 4-Domains A8 | 2.10 | 0.8 | 0.087 | 1.07E-64 |
| FOXJ1 | Forkhead Box J1 | 2.09 | 0.78 | 0.12 | 9.52E-46 |
| MORN5 | MORN Repeat Containing 5 | 2.08 | 0.78 | 0.08 | 3.05E-66 |
| SNTN | Sentan | 2.08 | 0.82 | 0.095 | 3.01E-58 |
| ODF3B | | 2.07 | 0.9 | 0.254 | 1.38E-30 |
| ZMYND10 | Zinc Finger MYND-Type Containing 10 | 2.05 | 0.8 | 0.081 | 5.07E-68 |
| CHI3L1 | Chitinase 3 Like 1 | 2.04 | 0.7 | 0.109 | 3.66E-41 |
| DYDC2 | DPY30 domain-containing protein 2 | 2.03 | 0.8 | 0.076 | 6.82E-73 |
| MMP1 | Matrix Metallopeptidase 1 | 2.03 | 0.54 | 0.027 | 1.45E-95 |
| C5orf49 | Cilia And Flagella Associated Protein 90 | 2.02 | 0.78 | 0.105 | 2.04E-48 |
| C9orf116 | Piercer Of Microtubule Wall 1 | 2.02 | 0.82 | 0.16 | 1.44E-36 |
| CSF2 | Colony Stimulating Factor 2 | 2.02 | 0.48 | 0.004 | 0 |
| HLA-DRA | Major Histocompatibility Complex, Class II, DR Alpha | 1.99 | 1 | 0.516 | 2.06E-22 |

#### Supplemental Figure Legends

**Figure S1.** FACS gating strategy showing selection of single endometrial cells in FSC area vs FSC (top left panel), followed by gating on EpCAM to separate viable $PI^{-}CD45^{-}$ endometrial cells into $EpCAM^{-}$ stromal/mesenchymal cells and $EpCAM^{+}$ epithelial cells (top middle panel). The $EpCAM^{-}$ stromal/mesenchymal cells were further gated and sorted into $SUSD2^{+}$ and $SUSD2^{-}$ subpopulations (lower left panel). $EpCAM^{+}$ epithelial cells were further gated into 4 subpopulations based on SSEA1 and N-cadherin (NCAD) (top left panel). The $EpCAM^{+}NCAD^{+}SSEA\text{-}1^{-}$ subpopulation was then sorted for high EpCAM expression (red arrow, lower right panel).

**Figure S2. Clusters identified in individual samples and initial cell type annotation.**

**(A)** UMAP plots showing contribution of individual samples to each cell cluster.

**(B)** UMAP plot of cells annotated to cell types using SingleR and the Human Cell Atlas (HCA) and the Blueprint and Encode as reference datasets. Cell types are listed with the cell number in the Table (left panel).

**Figure S3. Gene expression in endometrial cell populations and stem/progenitor markers in individual samples.**

**(A)** UMAP plot of *EPCAM* gene expression in each endometrial cell cluster.

**(B)** UMAP plot of *SOX9* gene expression in each endometrial cell cluster.

**(C)** UMAP plot of *STEAP4* gene expression in each endometrial cell cluster.

**(D)** UMAP plot of *FOXA2* gene expression in each endometrial cell cluster.

**(E)** UMAP plot of *AXIN2* gene expression in each endometrial cell cluster.

**(F)** UMAP plot of *SUSD2* gene expression in each endometrial cell cluster.

**(G)** UMAP plots *SUSD2* expression in each individual sample.

**(H)** UMAP plots of the expression of *CDH2* and *SOX9* combinations in individual samples.

**Figure S4. Gene ontology analysis of genes specific to the mesenchymal populations.**

**(A)** Biological processes enriched for genes highly expressed within the eMSC populations (Clusters 4 and 6).

**(B)** Biological processes enriched for genes highly expressed within the mature stromal populations including the decidual stromal cells (Cluster 0) and late secretory stroma (Cluster 8).

**Figure S5. Cell fate trajectory and cell cycle phase analysis.**

**(A)** RNA Velocity analysis showing unspliced–spliced phase portraits for *SUSD2.*

**(B)** RNA Velocity analysis showing unspliced–spliced phase portraits for *CDH2.*

**(C)** RNA Velocity analysis showing unspliced–spliced phase portraits for *SOX9.*

**(D)** Proportion of cells in each cluster at different phases of cell cycle: G1, S, G2/M. Cell population with cluster number in brackets.

**Figure S6. Gene ontology analysis of genes specific to the epithelial progenitor populations.** Biological processes enriched for genes highly expressed within the epithelial progenitor populations (Clusters 2 and 7).

**Figure S7. Steroid hormone receptor expression and associated co-regulators in endometrial cells.**

**(A)** *ESR1* and *PGR* and highly expressed in Epithelial Progenitor populations.

**(B)** Dot plot showing *ESR1, PGR* and steroid hormone co-activators and co-repressors in endometrial epithelial cell populations.

#### Supplemental Information

#### Supplemental Experimental Procedures

##### Cell-type annotation

Cells were annotated according to cell type using two supervised transcriptome-based cell-type classification methods. Previously published marker genes for endometrial cell types (Endometrial Epithelial Cells, *EPCAM, CDH1*; epithelial progenitors, *SOX9, AXIN2, CDH2*; glandular epithelial, *FOXA2*; ciliated epithelial, *FOXJ1*, *PIFO, TPPP3*; luminal epithelial, *LGR5, PTGS1, WNT7A*; *MUC5B*$^{+}$ epithelial, *MUC5B, TFF3, SAA1*; Endometrial Stromal Cells, *WT1, IGF1, PDGFRA*; Perivascular, *SUSD2, RGS5, NOTCH3* (Cousins et al., 2021; Fonseca et al., 2023; Garcia-Alonso et al., 2021; Lv et al., 2022; Tan et al., 2022; Wang et al., 2020) were used to assign clusters to cell types. Similarly, additional fine-scale subpopulations of stromal and *SOX9*$^{+}$ epithelial cells were annotated using marker genes published by

Garcia-Alonso et al. including basal fibroblasts (*C7, OGN, ACTA2*), non-decidualised proliferative stroma (*MMP11, CRABP2, ECM1*), decidualised secretory stroma (*IL15, CFD*), non-cycling *SOX9*$^{+}$*LGR5*$^{+}$ luminal epithelial cells (*SOX9, LGR5, KRT17, WNT7A*) and *SOX9*$^{+}$*LGR5*$^{-}$ basal gland epithelial cells (*SOX9, LGR5, IHH, EMID1*). The expression profile of each cell was also used to assign cell type annotations by comparing them to the Human Cell Atlas (HCA) and the Blueprint and Encode reference datasets using SingleR (v2.8.0) (Aran et al., 2019).

**Gene ontology analysis**

Gene enrichment and ontology analysis was performed using ShinyGO on the most significantly increased genes found in at least 75% of cells within clusters 2 and 7, and clusters 4, 6, 0 and 8.

**Immunofluorescence**

Immunofluorescence was performed on 8µm full thickness endometrium sections, previously fixed in 4% paraformaldehyde and cryopreserved in 30% sucrose and embedded in optimal cutting temperature compound (OCT). Unless stated all sections underwent the following protocol. Briefly, sections were thawed and rehydrated in PBS, and blocked in DAKO blocking solution (Agilent X0909) for 1 hour at room temperature (RT). Primary antibodies for Rabbit anti-N-cadherin (CST 3116 1/100), Mouse anti-SSEA-1 (Merck MAB4301 1/100), Rabbit anti-IHH (Abcam ab 52919 1/100), Rabbit anti-TRH (HPA033596 1/500) and Rabbit anti-BOC (Thermofisher BS1232R 1/100) were applied in 1%BSA:PBS overnight at 4°C. Rabbit IgG (Thermofisher 10500C) and Mouse IgM (Caltag MGM00) isotype controls were used at the same concentrations as primary antibodies. Sections were incubated with secondary antibodies donkey anti-rabbit AF488 (Thermofisher A21206 1/500), goat anti-mouse AF568 (Thermofisher A11004 1/500), or donkey anti rabbit AF647 (Abcam ab 150075 1/500) for 2hrs RT. In the case of double or triple immunostaining, antibodies were added sequentially and specific species blocks (20% goat serum:1%BSA:PBS or 20% donkey serum:1%BSA:PBS) were applied prior to the additional primary antibodies for 30mins RT. Nuclei were stained 61,with 5ug/ml Hoescht 33528 diluted in PBS for 5 mins RT. Slides were mounted in fluorescent mounting medium (DAKO S3203) and imaged using an Olympus FV1200 confocal microscope using a 20X objective lens. To obtain full thickness images, multiple fields of view were stitched together using the tile and stitch functions. Brightness and contrast were adjusted in a linear manner using FIJI software.

Aran, D., Looney, A.P., Liu, L., Wu, E., Fong, V., Hsu, A., Chak, S., Naikawadi, R.P., Wolters, P.J., Abate, A.R., et al. (2019). Reference-based analysis of lung single-cell sequencing reveals a transitional profibrotic macrophage. Nat. Immunol. *20*, 163-172. 10.1038/s41590-018-0276-y.

Cousins, F.L., Pandoy, R., Jin, S., and Gargett, C.E. (2021). The Elusive Endometrial Epithelial Stem/Progenitor Cells. Front Cell Dev Biol *9*, 640319. 10.3389/fcell.2021.640319.

Fonseca, M.A.S., Haro, M., Wright, K.N., Lin, X., Abbasi, F., Sun, J., Hernandez, L., Orr, N.L., Hong, J., Choi-Kuaea, Y., et al. (2023). Single-cell transcriptomic analysis of endometriosis. Nat. Genet. *55*, 255-267. 10.1038/s41588-022-01254-1.

Garcia-Alonso, L., Handfield, L.F., Roberts, K., Nikolakopoulou, K., Fernando, R.C., Gardner, L., Woodhams, B., Arutyunyan, A., Polanski, K., Hoo, R., et al. (2021). Mapping the temporal and spatial dynamics of the human endometrium in vivo and in vitro. Nat. Genet. *53*, 1698-1711. 10.1038/s41588-021-00972-2.

Lv, H., Zhao, G., Jiang, P., Wang, H., Wang, Z., Yao, S., Zhou, Z., Wang, L., Liu, D., Deng, W., et al. (2022). Deciphering the endometrial niche of human thin endometrium at single-cell resolution. Proc. Natl. Acad. Sci. U. S. A. *119*. 10.1073/pnas.2115912119.

Tan, Y., Flynn, W.F., Sivajothi, S., Luo, D., Bozal, S.B., Davé, M., Luciano, A.A., Robson, P., Luciano, D.E., and Courtois, E.T. (2022). Single-cell analysis of endometriosis reveals a coordinated transcriptional programme driving immunotolerance and angiogenesis across eutopic and ectopic tissues. Nat. Cell Biol. *24*, 1306-1318. 10.1038/s41556-022-00961-5.

Wang, W., Vilella, F., Alama, P., Moreno, I., Mignardi, M., Isakova, A., Pan, W., Simon, C., and Quake, S.R. (2020). Single-cell transcriptomic atlas of the human endometrium during the menstrual cycle. Nat. Med. *26*, 1644-1653. 10.1038/s41591-020-1040-z.

### Supplemental Figures

#### Figure S1

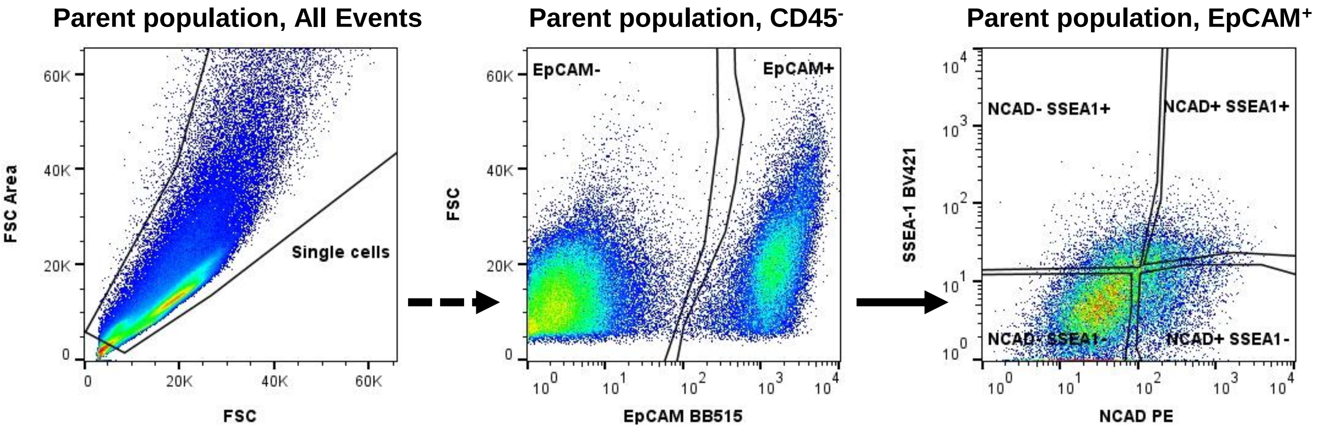


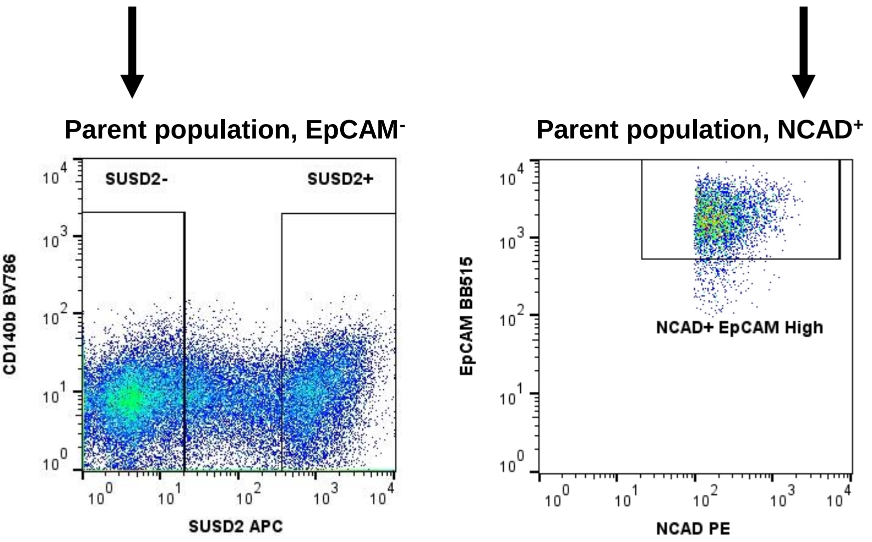


| Populations |
|---|
| All events |
| Single cells |
| Whole cells |
| Live cells |
| CD45⁻ |
| EpCAM⁺ |
| NCAD⁺SSEA-1⁻ |
| NCAD⁺SSEA-1⁺ |
| NCAD⁻SSEA-1⁺ |
| NCAD⁻SSEA-1⁻ |
| EpCAM⁻ |
| SUSD2⁻ |
| SUSD2⁺ |

Figure S2.

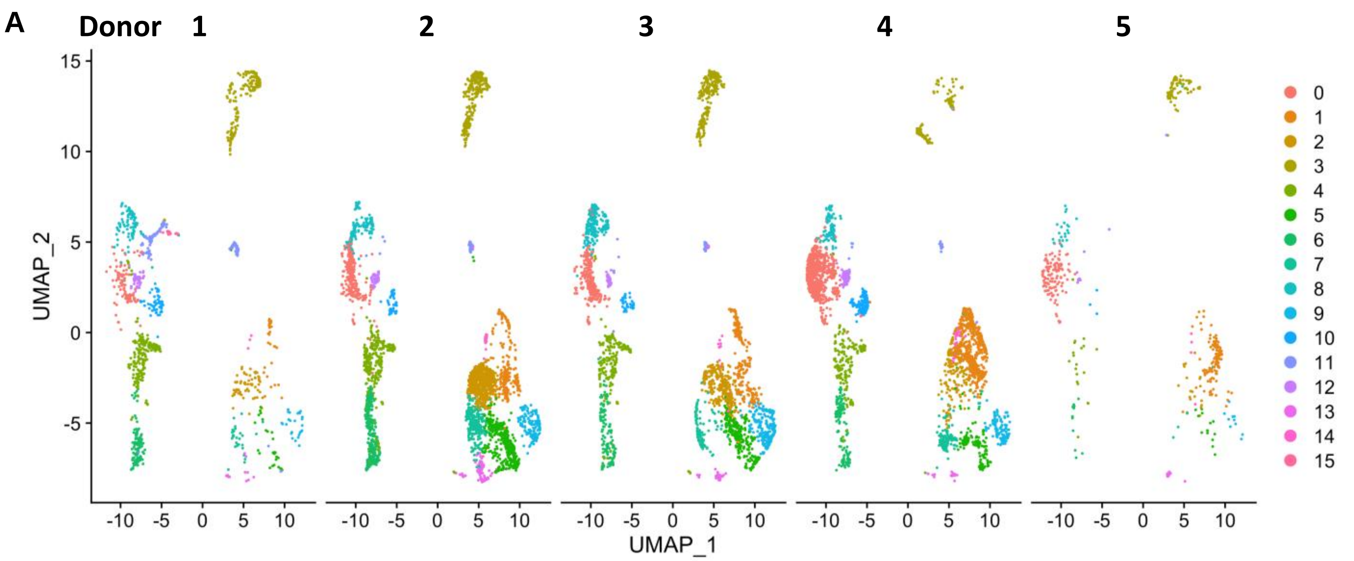


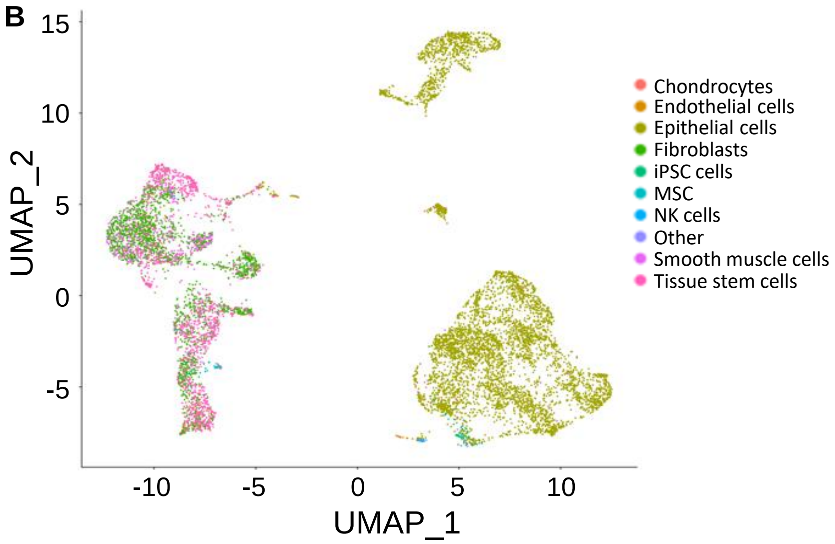


| Cell type | Cell No. |
|---|---|
| Epithelial | 4,462 |
| Tissue stem cells | 3,316 |
| Fibroblasts | 1,164 |
| Smooth muscle cells | 478 |
| iPSC | 70 |
| chondrocytes | 30 |
| MSC | 29 |
| NK cells | 15 |
| Endothelial cells | 13 |
| Neurons | 13 |

Figure S3.

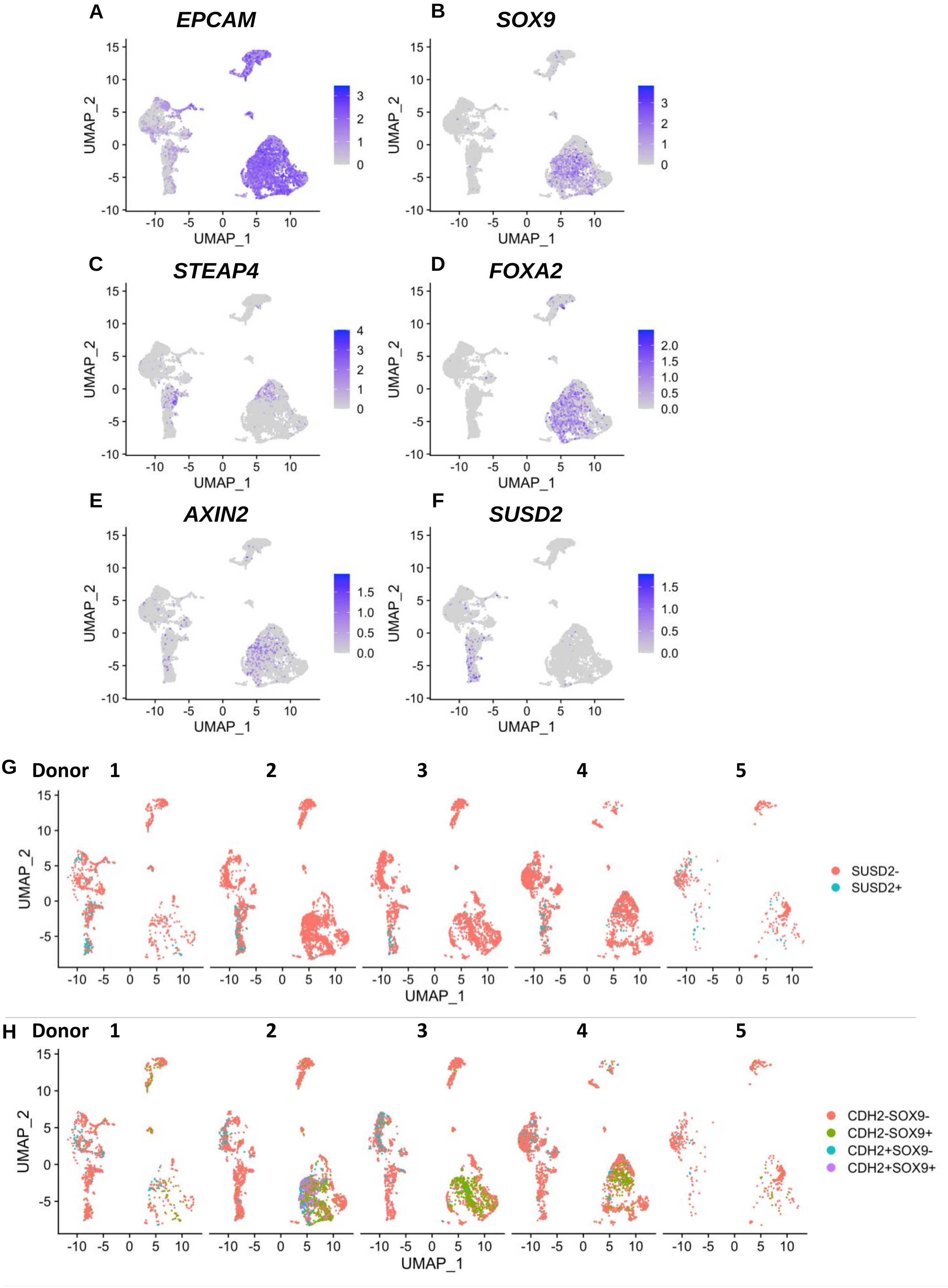

A
EPCAM
B
SOX9
C
STEAP4
D
FOXA2
E
AXIN2
F
SUSD2
UMAP_1
UMAP_2
G
Donor 1
2
3
4
5
SUSD2-
SUSD2+
H
Donor 1
2
3
4
5
CDH2-SOX9-
CDH2-SOX9+
CDH2+SOX9-
CDH2+SOX9+

Figure S4.

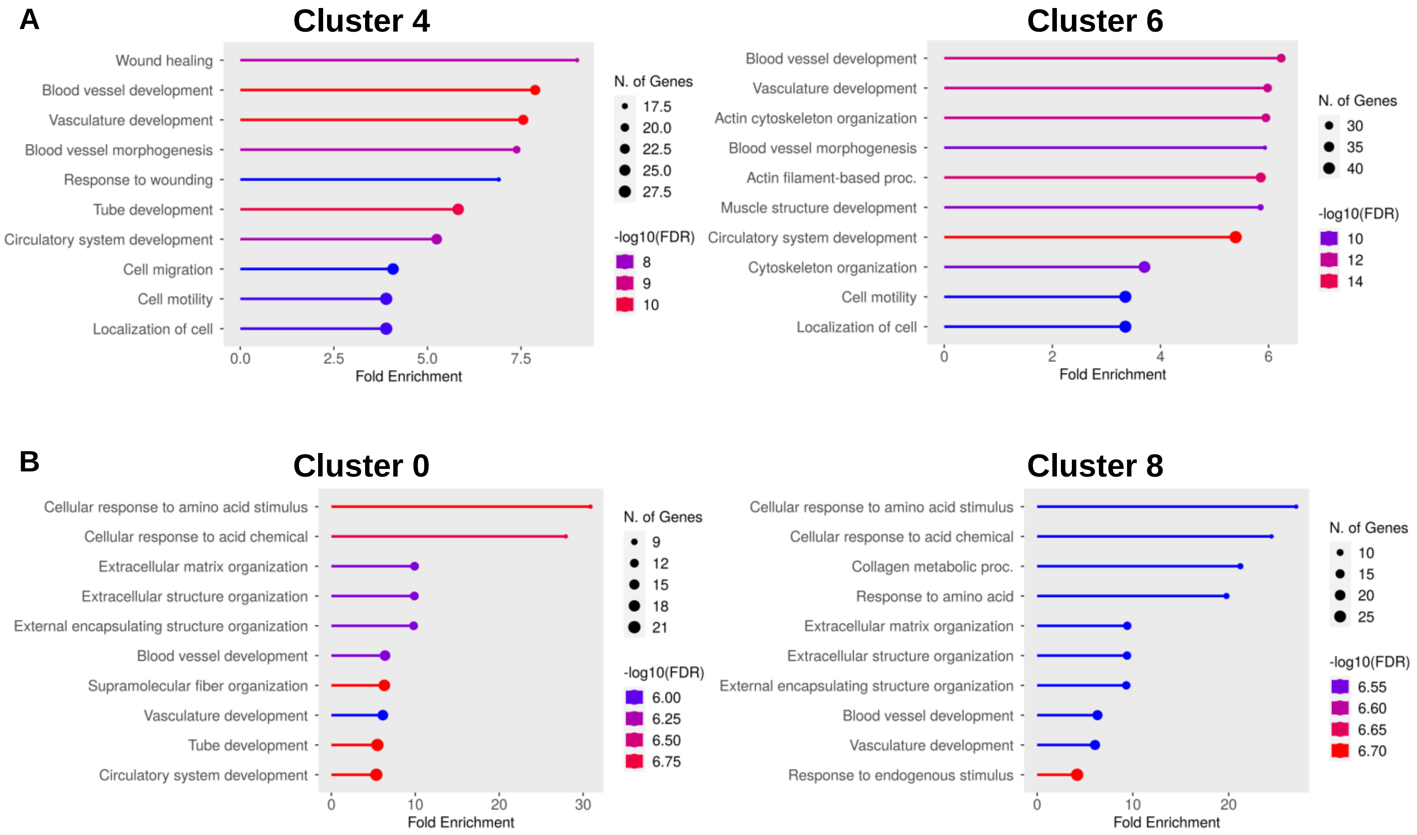

A
Cluster 4
Wound healing
Blood vessel development
Vasculature development
Blood vessel morphogenesis
Response to wounding
Tube development
Circulatory system development
Cell migration
Cell motility
Localization of cell
0.0
2.5
5.0
7.5
Fold Enrichment
N. of Genes
17.5
20.0
22.5
25.0
27.5
-log10(FDR)
8
9
10
Cluster 6
Blood vessel development
Vasculature development
Actin cytoskeleton organization
Blood vessel morphogenesis
Actin filament-based proc.
Muscle structure development
Circulatory system development
Cytoskeleton organization
Cell motility
Localization of cell
0
2
4
6
Fold Enrichment
N. of Genes
30
35
40
-log10(FDR)
10
12
14
B
Cluster 0
Cellular response to amino acid stimulus
Cellular response to acid chemical
Extracellular matrix organization
Extracellular structure organization
External encapsulating structure organization
Blood vessel development
Supramolecular fiber organization
Vasculature development
Tube development
Circulatory system development
0
10
20
30
Fold Enrichment
N. of Genes
9
12
15
18
21
-log10(FDR)
6.00
6.25
6.50
6.75
Cluster 8
Cellular response to amino acid stimulus
Cellular response to acid chemical
Collagen metabolic proc.
Response to amino acid
Extracellular matrix organization
Extracellular structure organization
External encapsulating structure organization
Blood vessel development
Vasculature development
Response to endogenous stimulus
0
10
20
Fold Enrichment
N. of Genes
10
15
20
25
-log10(FDR)
6.55
6.60
6.65
6.70

Figure S5.

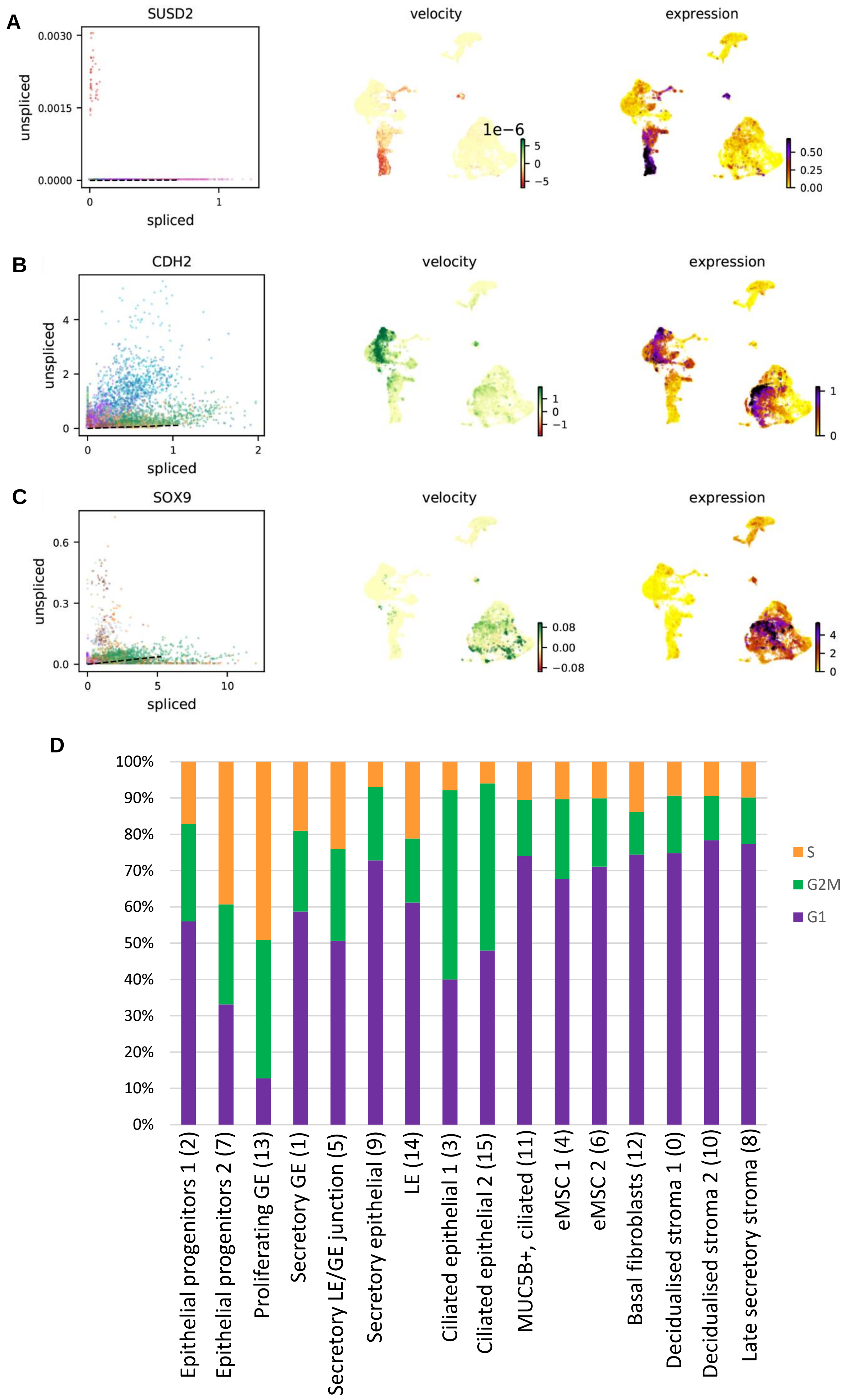

A
SUSD2
velocity
expression
unspliced
spliced
0.0030
0.0015
0.0000
0
1
1e−6
5
0
−5
0.50
0.25
0.00
B
CDH2
velocity
expression
unspliced
spliced
4
2
0
0
1
2
1
0
−1
1
0
C
SOX9
velocity
expression
unspliced
spliced
0.6
0.3
0.0
0
5
10
0.08
0.00
−0.08
4
2
0
D
100%
90%
80%
70%
60%
50%
40%
30%
20%
10%
0%
S
G2M
G1
Epithelial progenitors 1 (2)
Epithelial progenitors 2 (7)
Proliferating GE (13)
Secretory GE (1)
Secretory LE/GE junction (5)
Secretory epithelial (9)
LE (14)
Ciliated epithelial 1 (3)
Ciliated epithelial 2 (15)
MUC5B+, ciliated (11)
eMSC 1 (4)
eMSC 2 (6)
Basal fibroblasts (12)
Decidualised stroma 1 (0)
Decidualised stroma 2 (10)
Late secretory stroma (8)

### Figure S6.

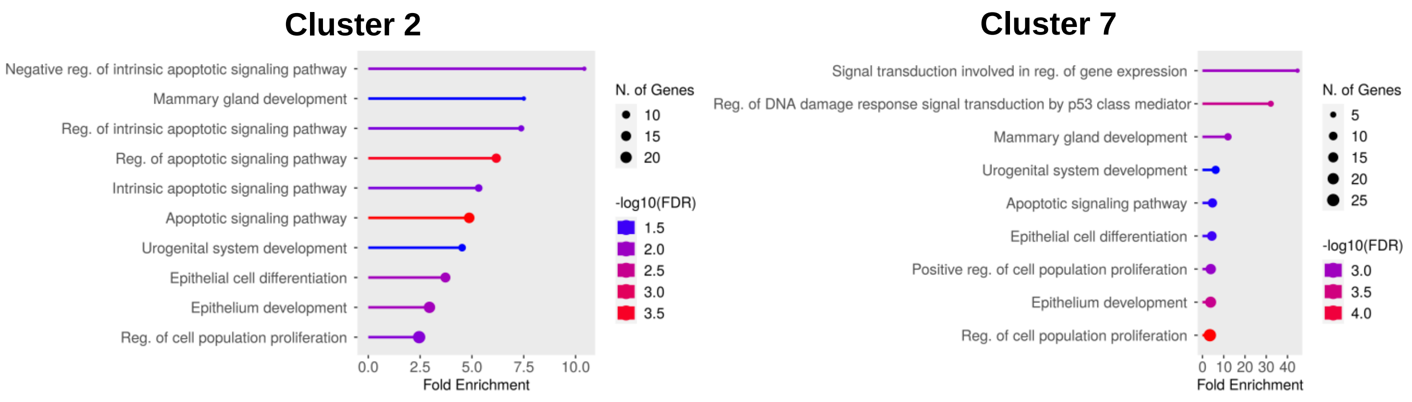
Cluster 2
Negative reg. of intrinsic apoptotic signaling pathway
Mammary gland development
Reg. of intrinsic apoptotic signaling pathway
Reg. of apoptotic signaling pathway
Intrinsic apoptotic signaling pathway
Apoptotic signaling pathway
Urogenital system development
Epithelial cell differentiation
Epithelium development
Reg. of cell population proliferation
0.0
2.5
5.0
7.5
10.0
Fold Enrichment
N. of Genes
10
15
20
-log10(FDR)
1.5
2.0
2.5
3.0
3.5
Cluster 7
Signal transduction involved in reg. of gene expression
Reg. of DNA damage response signal transduction by p53 class mediator
Mammary gland development
Urogenital system development
Apoptotic signaling pathway
Epithelial cell differentiation
Positive reg. of cell population proliferation
Epithelium development
Reg. of cell population proliferation
0
10
20
30
40
Fold Enrichment
N. of Genes
5
10
15
20
25
-log10(FDR)
3.0
3.5
4.0

Figure. S7

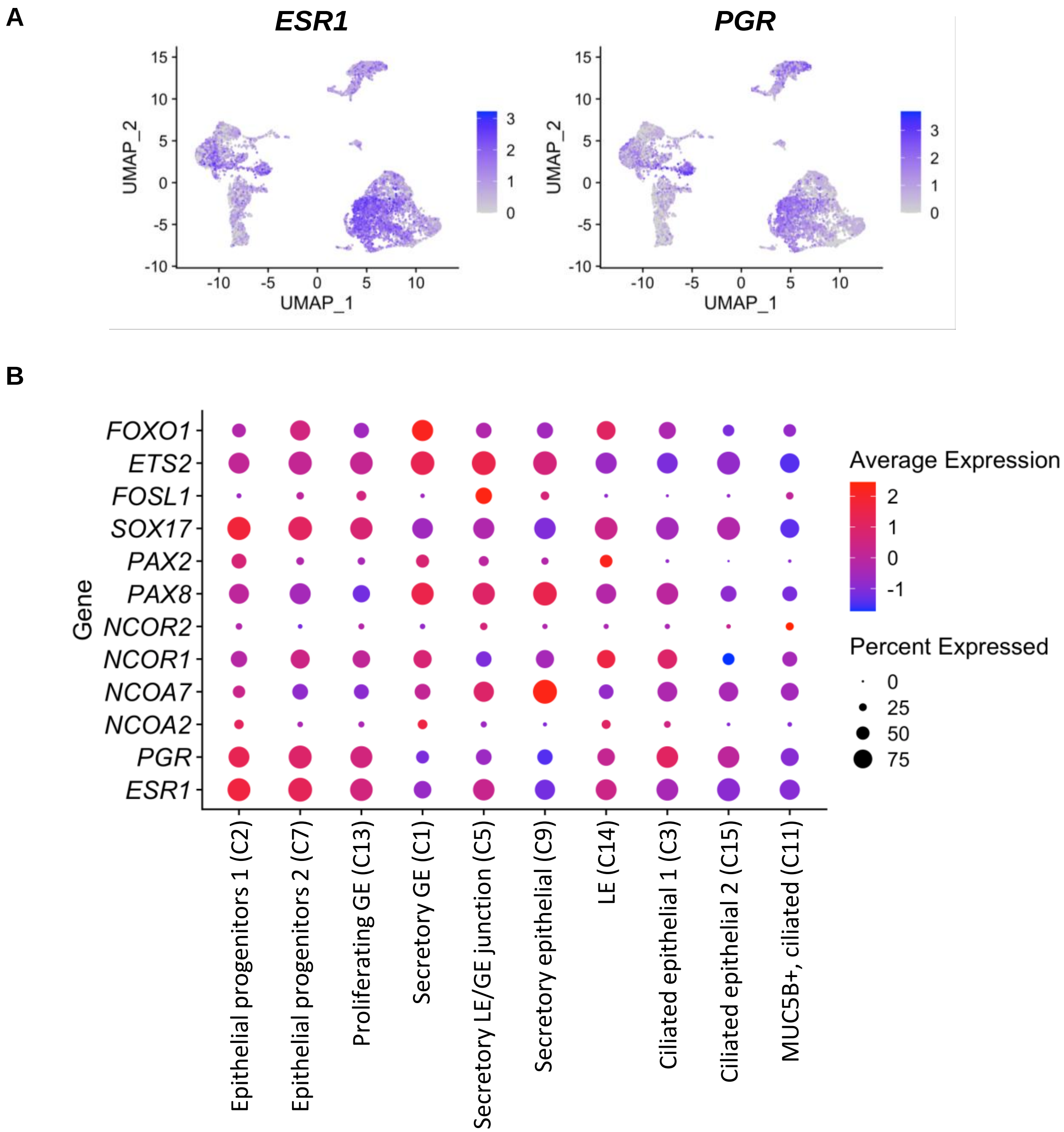
A
ESR1
PGR
UMAP_1
UMAP_2
B
Average Expression
Percent Expressed
Gene
FOXO1
ETS2
FOSL1
SOX17
PAX2
PAX8
NCOR2
NCOR1
NCOA7
NCOA2
PGR
ESR1
Epithelial progenitors 1 (C2)
Epithelial progenitors 2 (C7)
Proliferating GE (C13)
Secretory GE (C1)
Secretory LE/GE junction (C5)
Secretory epithelial (C9)
LE (C14)
Ciliated epithelial 1 (C3)
Ciliated epithelial 2 (C15)
MUC5B+, ciliated (C11)
